# Green Synthesis of Fluorescent Carbon Quantum Dots from Bearberry (Arctostaphylos uva-ursi) Extract via Hydrothermal and Microwave-Assisted Routes: Comparative Physicochemical Characterisation, Antioxidant Activity, and Biocompatibility Evaluation

**DOI:** 10.64898/2026.05.10.724067

**Authors:** Satyam Bhalerao, Jugal Patil, Abdul Mansuri, Ankur Singh, Bhagyesh Parmar, Ashutosh Kumar, Dhiraj Bhatia

## Abstract

Producing photoluminescent nanomaterials with controllable surface chemistry and predictable biological activity remains one of the outstanding problems in green nanoscience. The present study shows that, even when the same bearberry (*Arctostaphylos uva-ursi*) extract precursor is used, the mode of energy delivery during synthesis plays a determining role in shaping the surface composition, photophysical properties, and biological activity of the resulting carbon quantum dots (CQDs). Hydrothermal processing at 160 °C for 6 h yielded CQDs with an average particle size of 7.13 nm. Surface characterisation indicated abundant hydroxyl- and carbonyl-containing functionalities, while XPS analysis showed a comparatively higher proportion of graphitic sp^2^ carbon (43.06%). These structural features were accompanied by strong DPPH free-radical scavenging activity. Microwave-assisted synthesis, by contrast, yields 9.65 nm particles carrying a substantially greater surface carboxylate content (O–C=O: 19.06%), a higher fluorescence quantum yield, and enhanced intracellular uptake statistically significant in retinal epithelial cells at 200 µg/mL (p < 0.001) and showing concentration-dependent accumulation in zebrafish larvae from 100 µg/mL onwards (p < 0.05). XPS C 1s deconvolution, interpreted alongside FTIR difference spectroscopy, points to incomplete decarboxylation under microwave conditions as the primary mechanistic origin of these divergent properties. Cytocompatibility was uncompromised for both formulations across the full concentration range tested (10–250 µg/mL) in RPE-1 and HeLa cells, with no statistically significant loss of viability at any concentration. Taken together, these results define a synthesis-route-encoded structure–property relationship that permits rational selection between an antioxidant-optimised and an imaging-optimised CQD formulation from the same green precursor feedstock.

## 1. Introduction

Sub-10 nm in diameter and built around a graphitic carbon core, CQDs occupy an unusual position in the nanomaterials landscape: they offer the photoluminescent tunability of semiconductor quantum dots yet behave, in terms of toxicity and surface chemistry, much more like organic nanostructures^1,2^. Their existence was not anticipated - they were first noticed in 2004 as a fluorescent contaminant during electrophoretic purification of single-walled carbon nanotubes. Since then, the field has expanded considerably, and CQDs are now routinely employed in bioimaging, chemical sensing, photocatalysis, and drug delivery, applications made practical by their resistance to photobleaching, low cytotoxicity, and the relative ease with which their synthesis can be adapted to different precursor systems^3,4^. Photoluminescence in these materials spans the ultraviolet to near-infrared region and is typically excitation-dependent. This behaviour reflects at least three concurrent contributors: quantum confinement effects within sp^2^-hybridised graphitic domains, radiative recombination mediated by surface heteroatom functionalities, and molecular fluorophores generated in situ as carbonisation proceeds^5–8^.

What makes CQDs unusual among nanomaterials is how sensitively their optical and biological properties respond to changes in surface chemistry - and, by extension, to the choice of precursor and how it is processed^9^. Plant-derived precursors have attracted growing interest precisely because the native phytochemical matrix does much of the chemical work automatically. Polyphenols, organic acids, flavonoids, and nitrogen-bearing metabolites collectively enable carbonisation, surface passivation, and N/O co-doping within a single aqueous step, dispensing with the toxic reducing agents or harsh mineralising solvents that conventional routes often require^10,11^. Bearberry (*Arctostaphylos uva-ursi*), a circumpolar ericaceous shrub with a long history of medicinal use, presents a particularly well-suited precursor source. Its leaves are enriched in arbutin, hydroquinone, gallic acid, ellagic acid, and ursolic acid - compounds that carry phenolic hydroxyl and carbonyl groups positioned to undergo dehydration and cyclisation into polyaromatic carbon cores under thermal treatment^12^. Beyond their structural role in CQD formation, these phytochemicals are themselves potent radical scavengers, raising the question of whether CQDs synthesised from bearberry extract might inherit and perhaps amplify the antioxidant character of the source material.

The synthesis route represents a second, independent axis of control over CQD properties. In hydrothermal processing, organic precursors are held under autogenous pressure at elevated temperature for several hours; the result is a progressive, deep carbonisation that generates surfaces enriched in hydroxyl and carbonyl functionalities with a comparatively high graphitic carbon fraction^4,13^. Microwave-assisted synthesis operates through a fundamentally different mechanism - rapid, volumetric dielectric heating that drives nucleation within minutes but affords a much shorter effective reaction window, leaving a higher proportion of surface carboxylate groups intact and producing less graphitised cores^14^. Because these two routes differ only in how energy is delivered, applying both to an identical precursor formulation offers a controlled means of isolating the effect of thermal history on CQD structure and function. This approach is central to the present study.

In this study, CQDs were synthesised from bearberry extract co-processed with citric acid, urea, and PEG 2000 using both hydrothermal and microwave methods. The materials were comprehensively characterised in terms of their spectroscopic, compositional, and colloidal properties, and further assessed for biocompatibility, intracellular uptake, antioxidant activity, and in vivo fluorescence distribution in zebrafish larvae. The findings establish a consistent structure property relationship linking synthesis conditions to surface chemistry, optical behaviour, and biological performance, highlighting bearberry-derived CQDs as a promising photoluminescent, antioxidant-active nanomaterial platform.

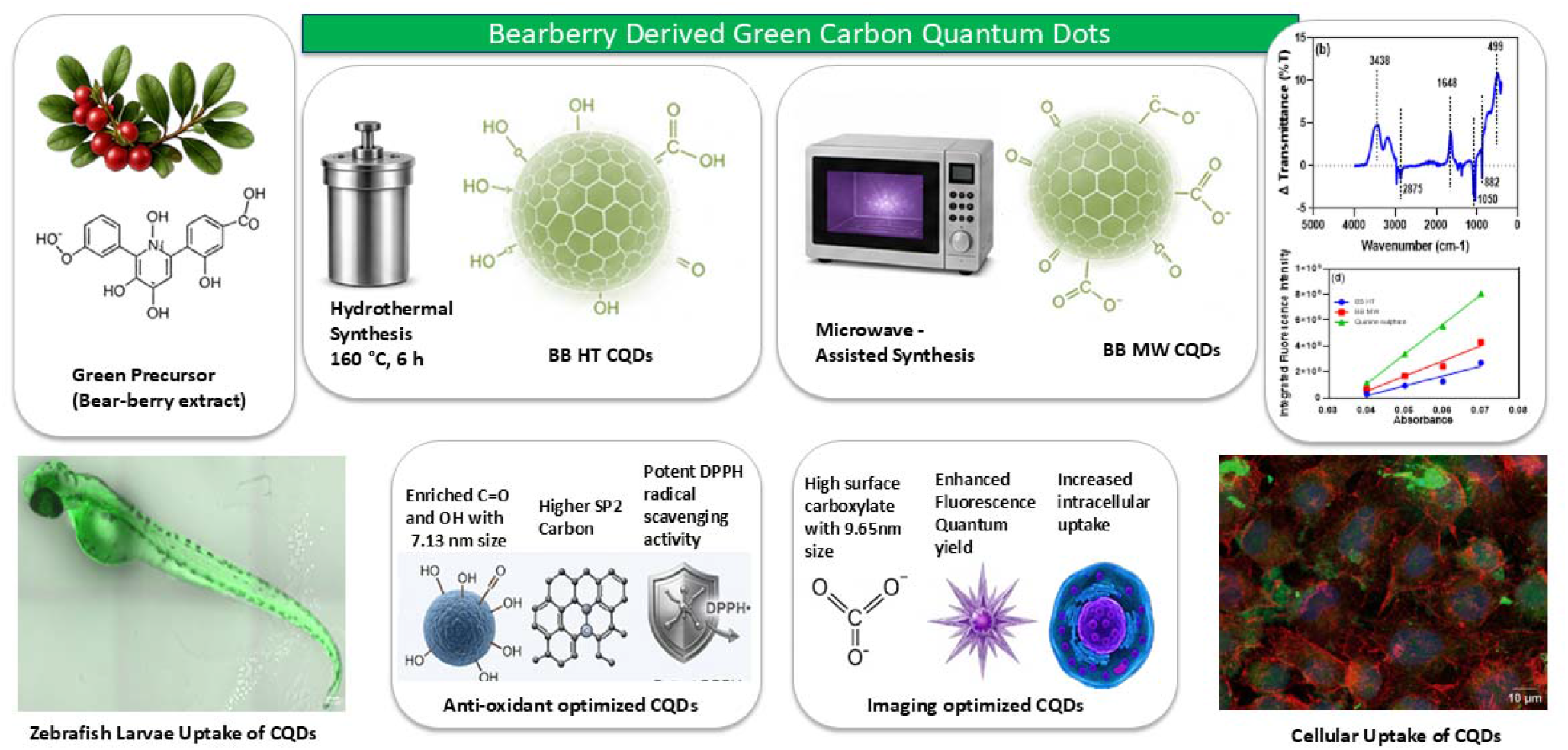

## 2. Materials and Methods

### 2.1 Materials

Bearberry (Arctostaphylos uva-ursi) herbal extract was sourced from BRL Chemicals (India). Citric acid (analytical grade, ≥99.5%), urea (analytical grade, ≥99%), and polyethylene glycol 2000 (PEG 2000, M□ ~2000 g/mol) were procured from commercial suppliers. Quinine sulphate dihydrate was used as a fluorescence quantum yield standard (QY = 0.54 in 0.1 M H□SO□ at 25 °C). 2,2-Diphenyl-1-picrylhydrazyl (DPPH) was employed to evaluate antioxidant activity. All aqueous preparations used ultrapure deionised water (resistivity ≥ 18.2 MΩ·cm). Culture media (DMEM), foetal bovine serum (FBS), and penicillin-streptomycin were of cell-culture grade. MTT stock solution [3-(4,5-dimethylthiazol-2-yl)-2,5-diphenyltetrazolium bromide] was prepared in phosphate-buffered saline (PBS) at a concentration of 5 mg/mL and subsequently filter-sterilised.

### 2.2 Hydrothermal Synthesis of BB HT CQDs

Bearberry extract was dissolved in 50% (v/v) aqueous ethanol at a 1:10 (w/v) ratio. Citric acid (0.5 g), urea (0.1 g), and PEG 2000 (50 mg) were added sequentially under magnetic stirring until complete dissolution. The resulting precursor solution was then loaded into a PTFE-lined stainless-steel autoclave. It was heated at 160 °C for 6 h in a muffle furnace. After natural cooling to room temperature, the reaction mixture was concentrated by rotary evaporation. The concentrate was filtered through a 0.22 µm polyethersulphone syringe filter to remove residual particulates. The filtrate was redissolved in deionised water and lyophilised for 48 h to obtain BB-HT CQDs as a dry powder.

### 2.3 Microwave-Assisted Synthesis of BB MW CQDs

The precursor composition was identical to that used in the hydrothermal route. This solution was poured into an open borosilicate glass beaker and irradiated at 750 W in a domestic microwave oven. Irradiation was sustained until the mixture had fully desiccated to a dry, charred residue. The residue was broken up and redispersed in deionised water, and the resulting suspension was concentrated under reduced pressure by rotary evaporation. Insoluble particulates were removed by passage through a 0.22 µm membrane filter, and the clarified filtrate was lyophilised for 48 h under conditions identical to those applied to BB-HT CQDs, affording BB-MW CQDs as a dry powder.

### 2.4 UV-Visible Absorption Spectroscopy

Absorption spectra were recorded by Thermo Scientific Evolution Pro UV-Vis Spectrophotometer from 200 to 800 nm against a deionised water blank. Sample optical densities were kept below 0.1 at the measurement wavelength to avoid inner-filter artefacts.

### 2.5 Photoluminescence Spectroscopy and Quantum Yield Determination

Steady-state emission spectra were acquired at a 350 nm excitation wavelength. Excitation-dependent spectra were recorded by varying the excitation from 222 to 450 nm in 50 nm increments. EEM maps were generated by scanning excitation from 250 to 450 nm and collecting emission from 300 to 700 nm. Fluorescence quantum yields were determined by the comparative gradient method using quinine sulphate as reference: QY□ = QY□ × (Slope□/Slope□) × (η□/η□)^2^, where Slope is the gradient of integrated emission intensity versus absorbance (five concentrations, OD < 0.07), and η is the solvent refractive index.

### 2.6 Fourier-Transform Infrared Spectroscopy

ATR-FTIR spectra of lyophilised powders were collected over 400-4000 cm□^1^ at 4 cm□^1^ resolution. A difference spectrum (ΔT = T(HT) − T(MW)) was computed after baseline correction to identify functional group enrichments specific to each synthesis route.

### 2.7 X-Ray Photoelectron Spectroscopy

XPS analysis was carried out using a monochromatic Al Kα radiation source (hν = 1486.6 eV). Survey scans were collected across a binding energy range of 0-1300 eV with a pass energy of 160 eV, while high-resolution C 1s spectra were obtained at a pass energy of 20 eV. All binding energies were referenced to adventitious C 1s at 284.8 eV. C 1s peaks were deconvoluted into five Gaussian components with constrained FWHM values to ensure physically meaningful fitting; goodness of fit was evaluated by R^2^.

### 2.8 MTT Cytotoxicity Assay

RPE-1 (non-cancerous human retinal pigment epithelium) and HeLa (cervical adenocarcinoma) cells were plated in 96-well plates at a density of 1 × 10□ cells per well and incubated for 24 h to enable proper adhesion. CQD suspensions prepared in complete DMEM were applied at concentrations of 10, 25, 50, 100, and 250 µg/mL, followed by incubation for 24 h at 37 °C in a humidified atmosphere containing 5% CO□. Subsequently, MTT reagent was introduced, the resulting formazan crystals were solubilised using DMSO, and absorbance was measured at 567 nm. Viability was expressed relative to untreated controls. Statistical evaluation was performed using one-way ANOVA followed by Dunnett’s post hoc multiple-comparison correction.

### 2.9 DPPH Radical Scavenging Assay

CQD dispersions at 25, 100, 250, and 1000 µg/mL were mixed with 0.1 mM DPPH solution prepared in ethanol and incubated for 30 min at room temperature under dark conditions. The absorbance was subsequently measured at 517 nm. Percentage inhibition = [(A_control − A_sample)/A_control] × 100. Ascorbic acid at identical concentrations served as a positive reference standard

### 2.10 In Vivo Fluorescence Uptake in Zebrafish Larvae

Wild-type zebrafish (Danio rerio) larvae at 72 h post-fertilisation were exposed to BB MW CQDs (50, 100, 250, and 500 µg/mL) in E3 for 4 h. Larvae were imaged using an laser scanning confocal microscope. Brightfield, fluorescence, and merged composite images were recorded under identical acquisition settings. Whole-larval normalised fluorescence intensity was quantified in ImageJ (using Raw intensity density/ area value) measurements corrected for background. Statistical analysis was carried out using one-way ANOVA followed by Dunnett’s post hoc test.

### 2.11 Dynamic Light Scattering and Zeta Potential

The hydrodynamic diameter distributions of both CQD formulations were determined by dynamic light scattering (DLS) at 25 °C using a Malvern Zetasizer Ultra instrument. Lyophilised powders were dissolved in ethanol (10mg/mL) and redispersed in deionised water at 0.1 mg mL□^1^, sonicated and filtered through a 0.22 µm membrane prior to analysis. Particle size is reported as the intensity-weighted mean hydrodynamic diameter (d.nm). Zeta potential measurements were performed using electrophoretic light scattering (ELS) on the same instrument with the corresponding dispersions.

### 2.12 Atomic Force Microscopy

Lyophilised CQD powders were reconstituted in deionised water at a concentration of 0.01 mg/mL and deposited onto freshly cleaved mica substrates by drop-casting, followed by vacuum drying at room temperature to remove residual solvent. AFM measurements were performed in tapping (intermittent-contact) mode. Two-dimensional height maps and three-dimensional topographic renders were processed using JPKSPM data processing software. Peak height values were extracted from the instrument colour scale of each height map. Particle surface density was assessed by visual inspection of the 2D topography images over multiple independently acquired scan areas.

### 2.13 Confocal Fluorescence Microscopy of Cellular Uptake

RPE-1 and HeLa cells were plated on glass coverslips in 24-well plates at a seeding density of 1 × 10□ cells per well and incubated for 24 h in complete DMEM to permit cell attachment. CQD dispersions were added at final concentrations of 100, 200, and 300 µg mL□^1^ and incubated for 20 min at 37 °C under 5% CO□. After incubation, the cells were rinsed three times with ice-cold PBS to eliminate residual unbound material. Fixation was carried out using 4% paraformaldehyde for 15 min at ambient temperature, followed by permeabilization with 0.1% Triton X-100 prepared in PBS for an additional 15 min. F-actin filaments were labelled with phalloidin–FITC (green fluorescence), while cell nuclei were stained with DAPI (blue fluorescence). CQD fluorescence was imaged directly using excitation at 405 nm, with emission collected in the 450-550 nm range. Imaging was performed on a Super-resolution microscope using identical acquisition settings across all treatment groups. Intracellular fluorescence was quantified in ImageJ using raw integrated density normalised to cell area. Values were background-corrected and expressed relative to the untreated control. Statistical analysis was performed using one-way ANOVA with Dunnett’s post hoc test. Significance thresholds were defined as # (p < 0.1) and *** (p < 0.001).***

## 3. Results and Discussion

### 3.1 Synthesis Strategy and Mechanistic Rationale

Bearberry extract, citric acid, urea, and PEG-2000 were chosen as the four components of the precursor system in order to take advantage of each component’s complementary roles in the synthesis of CQD. Bearberry extract acts as a phytochemically rich carbon source. It contains arbutin, gallic acid, and ellagic acid. These molecules are enriched in phenolic hydroxyl and carboxyl groups. Such functionalities promote condensation into polyaromatic domains during thermal treatment^15^. Citric acid functions as a classical carbon precursor. It undergoes dehydration and cyclisation to generate six-membered aromatic structures. These intermediates act as nucleation centres for graphitic core formation^16^. Urea provides a source of nitrogen doping. It introduces pyridinic and pyrrolic nitrogen species into the carbon framework and surface. This modifies both the electronic structure and surface reactivity of the resulting CQDs^16^. PEG 2000 acts as a passivating agent on the surface. It improves aqueous dispersibility, colloidal stability, and interparticle aggregation^17^.

The temperature profiles of the hydrothermal and microwave methods are essentially different. In the sealed autoclave, autogenous pressure develops as the solvent is heated above its boiling point. Sustained heating at 160 °C for 6 h promotes extensive dehydration, condensation, and progressive graphitisation of the organic precursors. In contrast, microwave irradiation at 750 W induces rapid volumetric heating via dielectric coupling with polar species. Nucleation is initiated within minutes. However, the open-vessel configuration and short reaction time limit the extent of carbonisation and structural ordering. These differences in reaction dynamics lead to distinct chemical outcomes. The surface functional group distributions are covered in the next sections (**Figure 1**).

**Figure 1.**
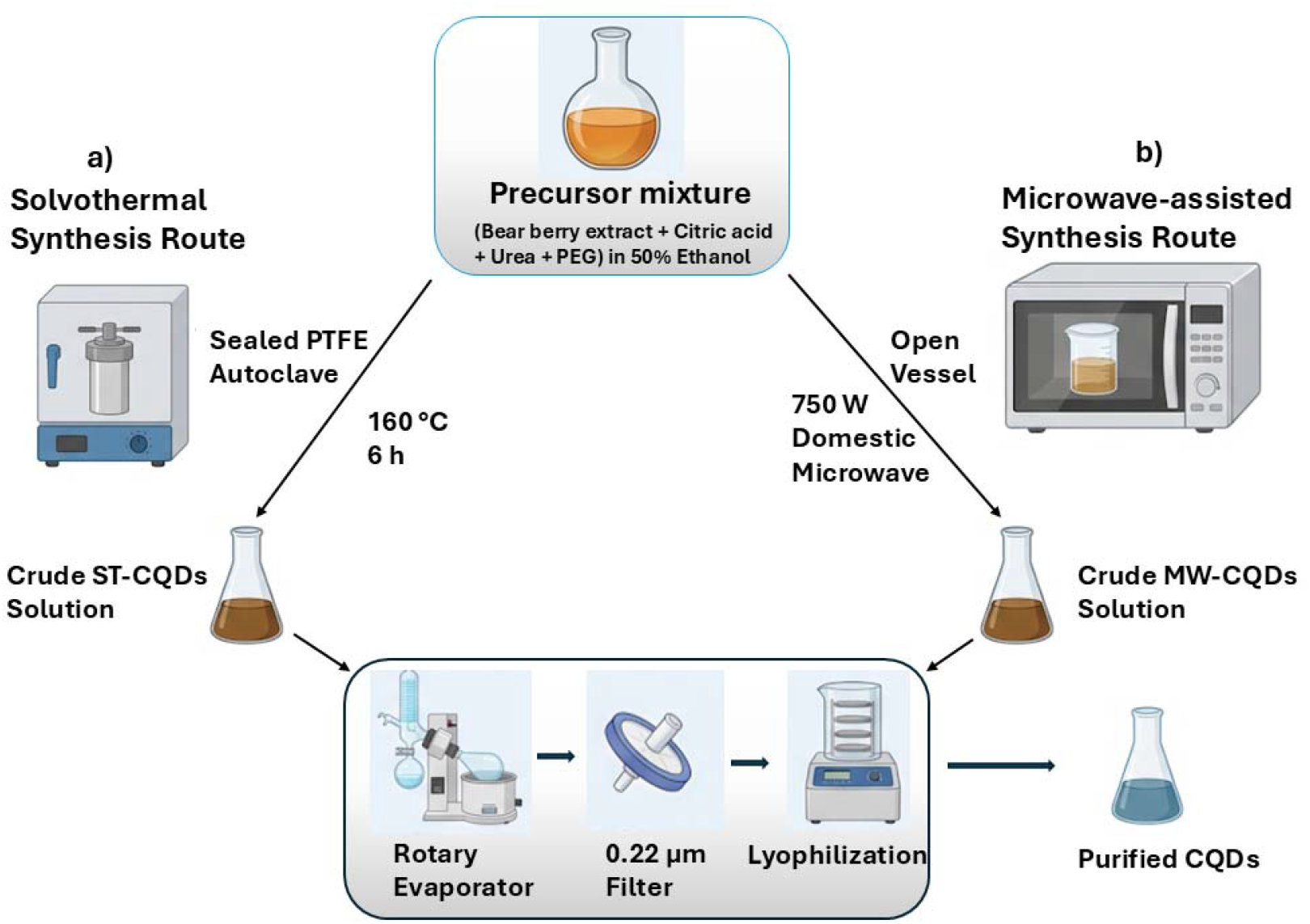
Schematic illustration of the green synthesis of bearberry-derived CQDs using (a) hydrothermal treatment (160 °C, 6 h, sealed PTFE autoclave) and (b) microwave irradiation (750 W domestic microwave). Both methods employed bearberry extract with citric acid, urea, and PEG-2000 in 50% aqueous ethanol, followed by identical purification steps including rotary evaporation, 0.22 µm filtration, and 48 h lyophilisation.

### 3.2 UV-Visible Absorption and Photoluminescence Characteristics

The UV-visible absorption spectra of both BB HT and BB MW CQDs (**Figure 2**) display two characteristic transitions consistent with graphitic CQD architecture. A dominant absorption near 210 nm corresponds to the π-π* transition of sp^2^-hybridised C=C bonds within the graphitic core. A secondary, bathochromically shifted absorption feature appearing at ~340 nm for BB-MW-CNPs and at a slightly red-shifted ~360 nm for BB-HT-CNPs corresponds to the n-π* transition arising from lone-pair electrons of surface carbonyl (C=O), carboxylate (O-C=O), and potentially imine (C=N) functional groups. The greater prominence and red-shift of the n-π* band in BB-HT-CNPs is consistent with the longer carbonisation time and higher thermal energy of the hydrothermal route, which promotes more extensive dehydration-condensation reactions and thus a denser population of oxygen-containing surface defect states, as subsequently confirmed by FTIR and XPS. A summary of the photophysical properties of both preparations is provided in **Table 1**.

**Figure 2.**
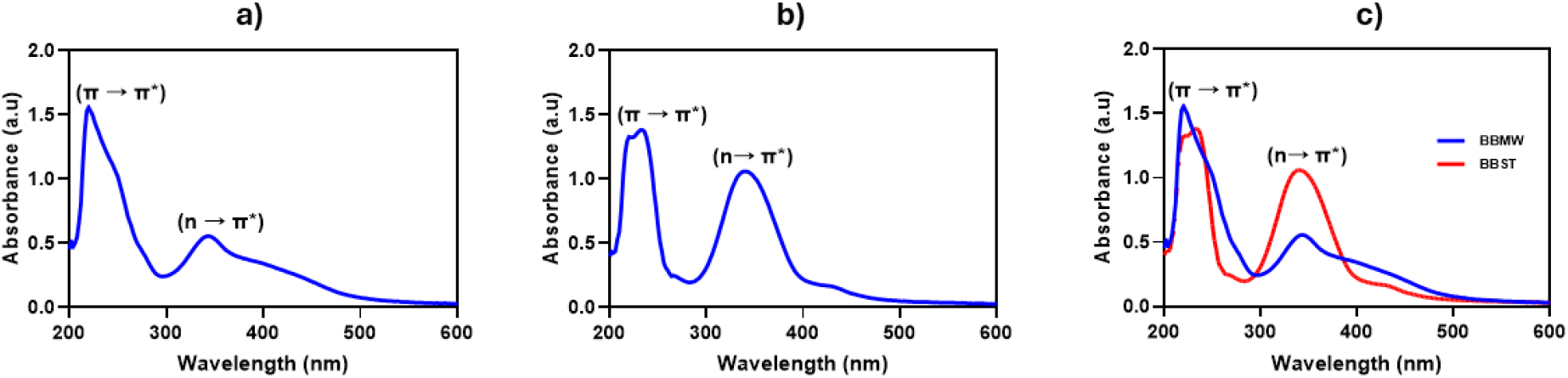
UV–visible absorption spectra of (a) BB MW CQDs and (b) BB HT CQDs recorded in dilute aqueous dispersion (optical density < 0.1 at the measurement wavelength). (c) Overlaid comparison of both spectra, highlighting the π-π* electronic transition of graphitic C=C domains (~210 nm) and the n-π* transition of surface carbonyl/carboxyl groups (~340 nm). The pronounced n-π* feature in BB HT CQDs relative to the broad shoulder observed for BB MW CQDs is consistent with the greater surface carbonyl density of the hydrothermally synthesised material.

**Table 1.** Summary of photophysical properties of BB HT CQDs and BB MW CQDs.

| Property | BB HT CQDs | BB MW CQDs |
| --- | --- | --- |
| $\pi \rightarrow \pi^*$ absorption maximum (nm) | ~210 | ~210 |
| $n \rightarrow \pi^*$ absorption feature | ~340 nm (distinct band) | ~340 nm (broad shoulder) |
| Emission maximum, $\lambda_{ex} = 350$ nm (nm) | 478 | 467 |
| Stokes shift (nm, $\lambda_{ex} = 350$ nm) | ~128 | ~117 |
| EEM dominant peak: $\lambda_{ex} / \lambda_{em}$ (nm) | 351 / 481 | 351 / 469 |
| Relative quantum yield (vs. quinine sulphate) | Lower | Higher |
| $n \rightarrow \pi^*$ band prominence | High (distinct peak) | Moderate (broad shoulder) |

Under 350 nm excitation, BB MW CQDs emit at a maximum of 467 nm whilst BB HT CQDs show a bathochromically shifted maximum at 478 nm (**Figure 3**). This 11 nm red-shift in the hydrothermal product is attributable to the greater density of surface defect states principally carbonyl and hydroxyl groups which introduce additional sub-band energy levels facilitating lower-energy radiative recombination^18^. The Stokes shifts are correspondingly larger in BB HT CQDs (~128 nm) than in BB MW CQDs (~117 nm), again consistent with a broader and lower-energy distribution of emissive surface states in the more oxidised material.

**Figure 3.**
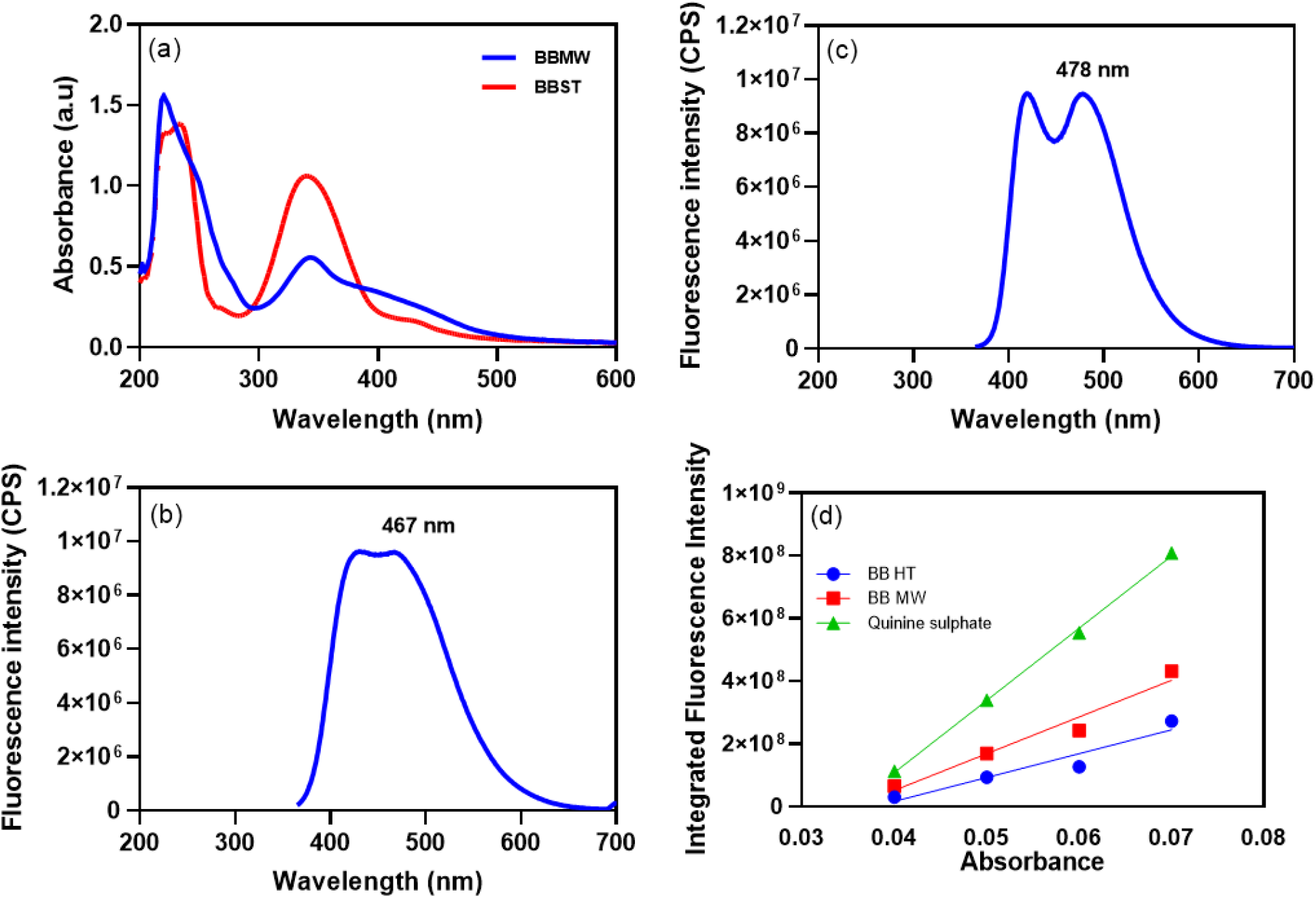
Photoluminescence emission spectra and quantum yield determination of BB CQDs. (a) Overlaid emission spectra of BB HT CQDs (red) and BB MW CQDs (blue) acquired at an excitation wavelength of 350 nm, highlighting the 11 nm bathochromic shift in the hydrothermally derived material (emission maxima at 478 nm and 467 nm, respectively). (b) Emission spectrum of BB MW CQDs (λex = 350 nm; peak at 467 nm). (c) Emission spectrum of BB HT CQDs (λex = 350 nm; peak at 478 nm). (d) Fluorescence quantum yield determination by the comparative method using quinine sulphate (QY = 0.54 in 0.1 M H□SO□) as reference; integrated fluorescence intensity is plotted against absorbance for BB HT CQDs (blue circles), BB MW CQDs (red squares), and quinine sulphate (green triangles). Lines represent linear regression fits; the slope ratio method was applied to calculate quantum yields.

Quantum yields were determined by the linear regression-based comparative method against quinine sulphate (QY = 0.54 in 0.1 M H□SO□). The relative slope analysis (**Figure 3d**) indicates that BB MW CQDs achieve a higher QY than BB HT CQDs a relationship mechanistically linked to the lower surface defect density of the microwave product. In CQDs, surface oxidative defects that confer antioxidant activity simultaneously introduce non-radiative recombination pathways that diminish quantum yield; the MW synthesis, by generating fewer such sites, thus produces particles that are less potent antioxidants but more efficient fluorophores. This antioxidant-fluorescence trade-off is an intrinsic consequence of surface chemistry and is a recurring motif in the CQD literature^19^.

Excitation-dependent photoluminescence spectra (**Figure 4**) confirm that both CQDs preparations exhibit the multicentre emission characteristic of surface-state heterogeneity. In both materials, emission intensity peaks at a 350 nm excitation wavelength and the emission maximum red-shifts progressively as the excitation wavelength increases behaviour arising from preferential excitation of increasingly lower-energy surface states as excitation photon energy decreases^20^. The EEM contour maps quantify this behaviour: the dominant fluorescence maximum for BB HT CQDs lies at λex/λem = 351/481 nm, versus 351/469 nm for BB MW CQDs, a 12 nm shift in the EEM peak that mirrors the single-wavelength emission data and corroborates the consistent influence of surface oxidation on emission energy across the entire excitation landscape.

**Figure 4.**
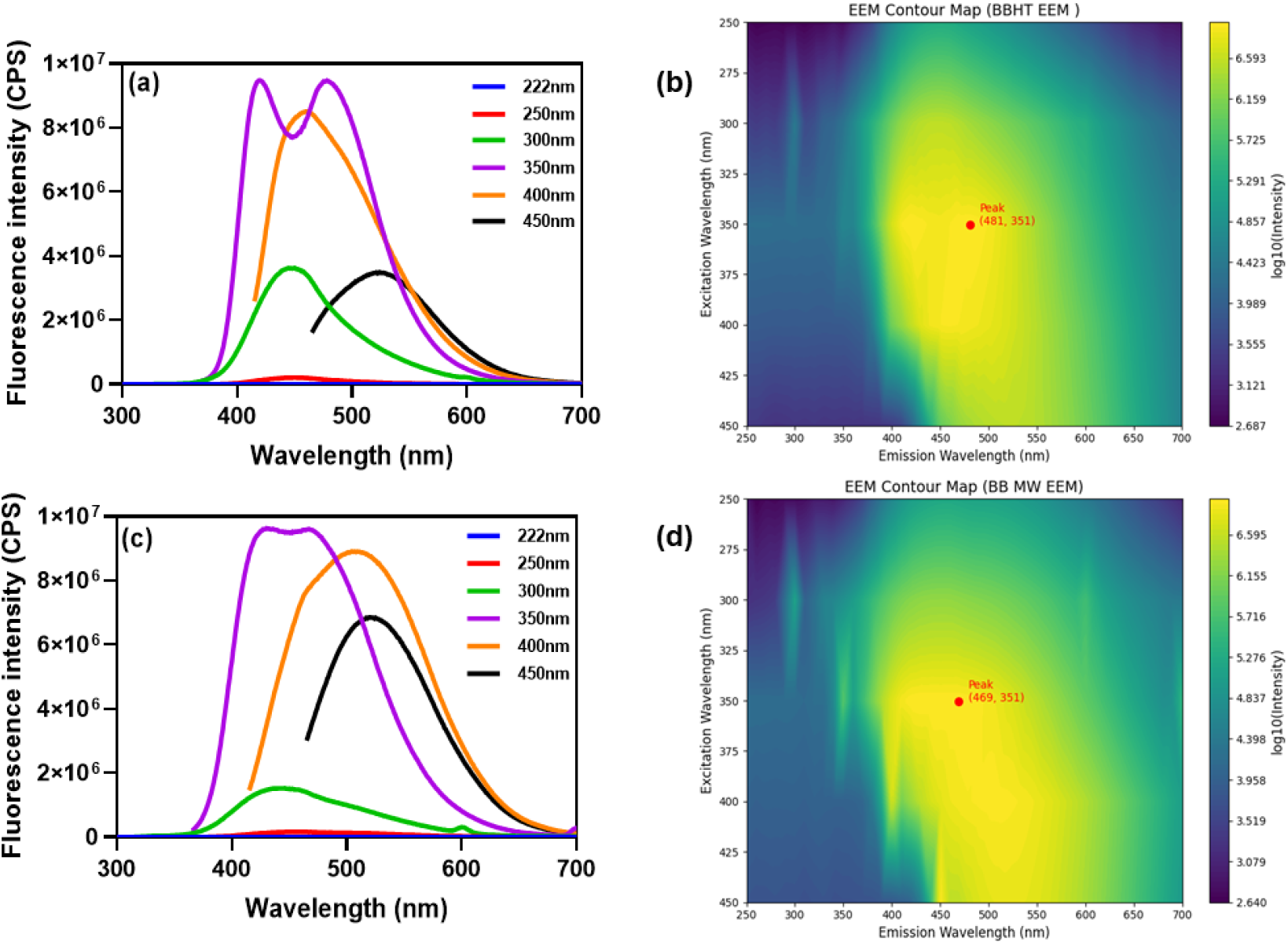
Excitation-dependent photoluminescence behaviour and excitation–emission matrix (EEM) fluorescence mapping of BB CQDs. Emission spectra of (a) BB HT CQDs and (c) BB MW CQDs collected at excitation wavelengths of 222, 250, 300, 350, 400, and 450 nm, demonstrating characteristic excitation-dependent fluorescence behaviour arising from the heterogeneous distribution of surface emissive states. EEM fluorescence contour maps (log□□ intensity scale) of (b) BB HT CQDs and (d) BB MW CQDs; red markers indicate the dominant fluorescence maxima at λex/λem = 351/481 nm and 351/469 nm, respectively. The 12 nm red shift in the EEM emission peak of BB HT CQDs relative to BB MW CQDs is consistent with greater surface oxidation and defect-state-mediated radiative recombination in the hydrothermally derived material.

### 3.3 FTIR Spectroscopic Characterisation

The ATR-FTIR spectra of BB-HT and BB-MW CQDs **(Figure 5**) display a similar set of absorption bands. This indicates that both synthesis routes yield carbon nanomaterials with comparable surface functionalities. A broad and intense band centred around 3400 cm□^1^ is observed in both samples. This band is the result of the joint contribution of N–H stretching modes from amine and amide groups created by urea-mediated nitrogen incorporation and O–H stretching vibrations linked to surface hydroxyl functionalities and adsorbed water molecules. Near 2900 cm□^1^, aliphatic C-H stretching bands emerge. Remaining −CH□- and −CH□ groups are responsible for these. There is complexity in the area between 1600 and 1650 cm□^1^. It includes contributions from amide II bending modes, C=C stretching of sp^2^-conjugated domains, and C=O stretching (carbonyl and amide I). The presence of ether and alcohol functionalities is confirmed by a high absorption about 1050 cm□^1^, which corresponds to C-O and C-O-C stretching. Additionally, a band at 880 cm□^1^ that is ascribed to aromatic C-H out-of-plane bending indicates peripheral aromatic structures.

**Figure 5.**
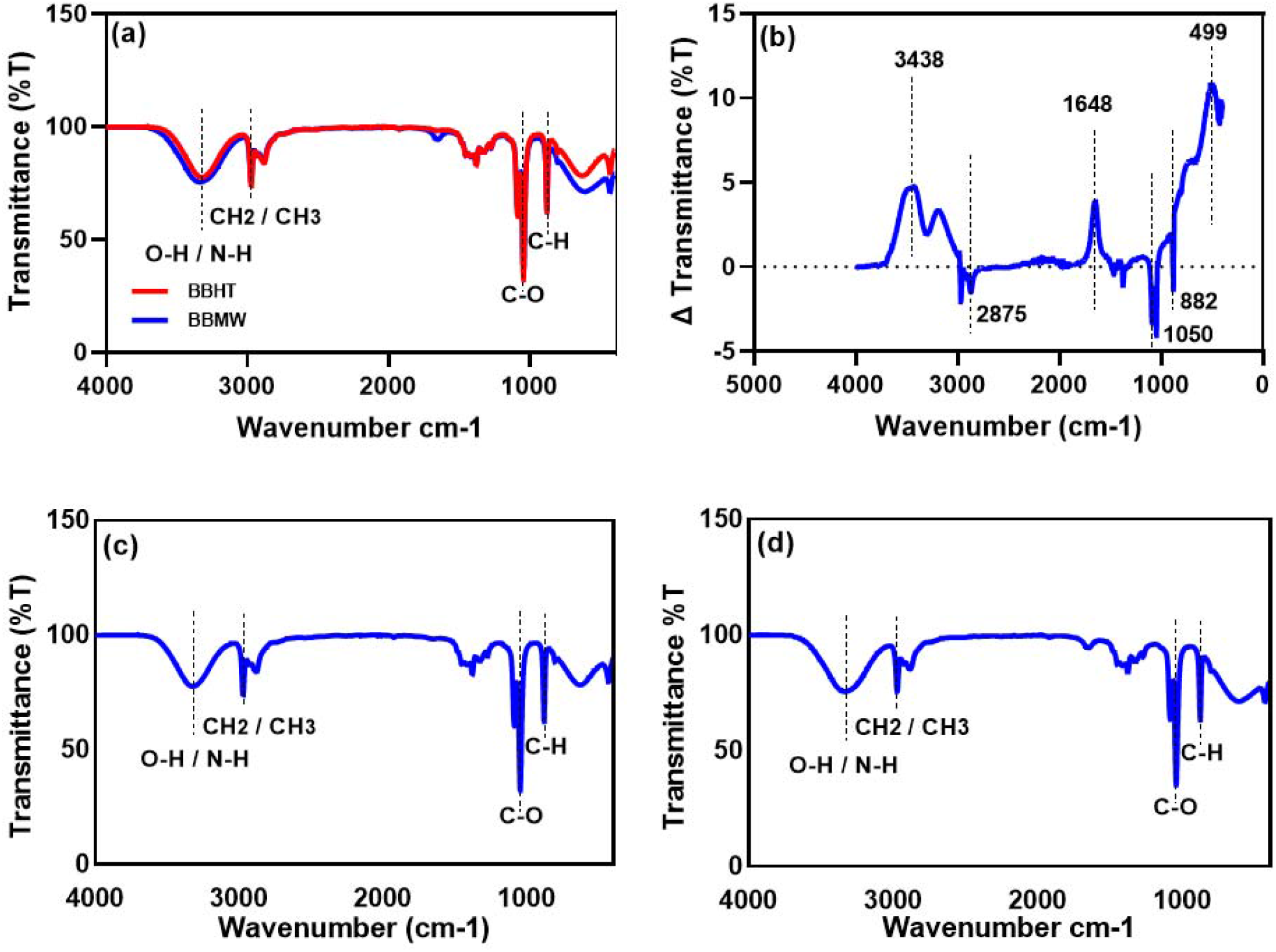
FTIR spectroscopic characterisation of BB CQDs. (a) Overlaid ATR-FTIR spectra of BB HT CQDs (red) and BB MW CQDs (blue) with shared vertical reference lines at key wavenumber positions. (b) Difference spectrum (ΔT = T(HT) − T(MW)); positive features indicate greater abundance in BB HT CQDs and negative features indicate enrichment in BB MW CQDs; dashed horizontal line at ΔT = 0 demarcates the two groups. Individual ATR-FTIR spectra of (c) BB HT CQDs and (d) BB MW CQDs. Assignments: 3438 cm□^1^ (O-H/N-H stretch), 2875 cm□^1^ (aliphatic C-H stretch), 1648 cm□^1^ (C=O/C=C/amide), 1050 cm□^1^ (C-O/C-O-C stretch), 882 cm□^1^ (aromatic C-H out-of-plane bend), 499 cm□^1^ (skeletal C-C/ring vibrations). See Table 2 for detailed functional group assignments and mechanistic interpretation.

The route-dependent divergence in surface chemistry is directly and quantitatively demonstrated by the difference spectrum (ΔT = T(HT) − T(MW)) (**Figure 5b**), which is explained in full in **Table 2**. Positive features in the difference spectrum at 3438, 1648, and 499 cm□^1^ establish that BB HT CQDs contain a higher abundance of O-H/N-H groups, carbonyl/conjugated functionalities, and ordered skeletal vibrations, respectively. Negative features at 2875, 1050, and 882 cm□^1^ indicate that BB MW CQDs are enriched in aliphatic C-H, ether/alcohol C-O, and aromatic C-H environments relative to the HT product^21^. This pattern is mechanistically coherent: the prolonged hydrothermal treatment at 160 °C progressively oxidises aliphatic domains and aromatic edge carbons, converting sp^3^ centres and peripheral aromatic sites into surface oxygenated functionalities, while the shorter microwave process preserves a comparatively higher proportion of these partially carbonised structures^22^.

**Table 2.**
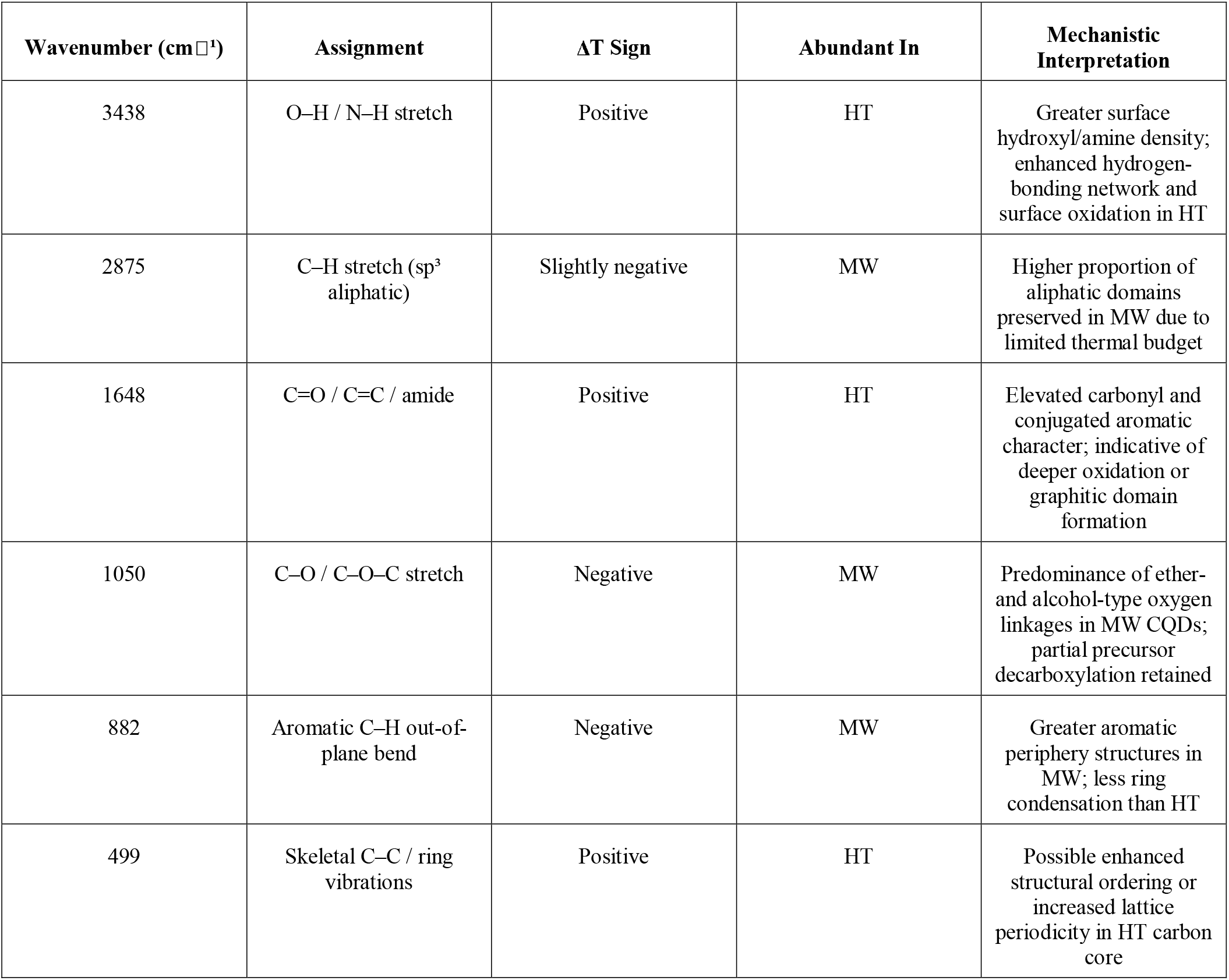
Comparative FTIR functional group analysis derived from the difference spectrum (BB HT - BB MW). Positive ΔT indicates enrichment in BB HT CQDs; negative ΔT indicates enrichment in BB MW CQDs.

| Wavenumber ($\text{cm}^{-1}$) | Assignment | $\Delta T$ Sign | Abundant In | Mechanistic Interpretation |
| --- | --- | --- | --- | --- |
| 3438 | O-H / N-H stretch | Positive | HT | Greater surface hydroxyl/amine density; enhanced hydrogen-bonding network and surface oxidation in HT |
| 2875 | C-H stretch ($\text{sp}^3$ aliphatic) | Slightly negative | MW | Higher proportion of aliphatic domains preserved in MW due to limited thermal budget |
| 1648 | C=O / C=C / amide | Positive | HT | Elevated carbonyl and conjugated aromatic character; indicative of deeper oxidation or graphitic domain formation |
| 1050 | C-O / C-O-C stretch | Negative | MW | Predominance of ether- and alcohol-type oxygen linkages in MW CQDs; partial precursor decarboxylation retained |
| 882 | Aromatic C-H out-of-plane bend | Negative | MW | Greater aromatic periphery structures in MW; less ring condensation than HT |
| 499 | Skeletal C-C / ring vibrations | Positive | HT | Possible enhanced structural ordering or increased lattice periodicity in HT carbon core |

### 3.4 X-Ray Photoelectron Spectroscopic Analysis

XPS survey spectra (**Figure 6a, b**) confirm the tri-elemental C/N/O composition of both preparations. Elemental abundances and derived atomic ratios are summarised in **Table 3**.

**Figure 6.**
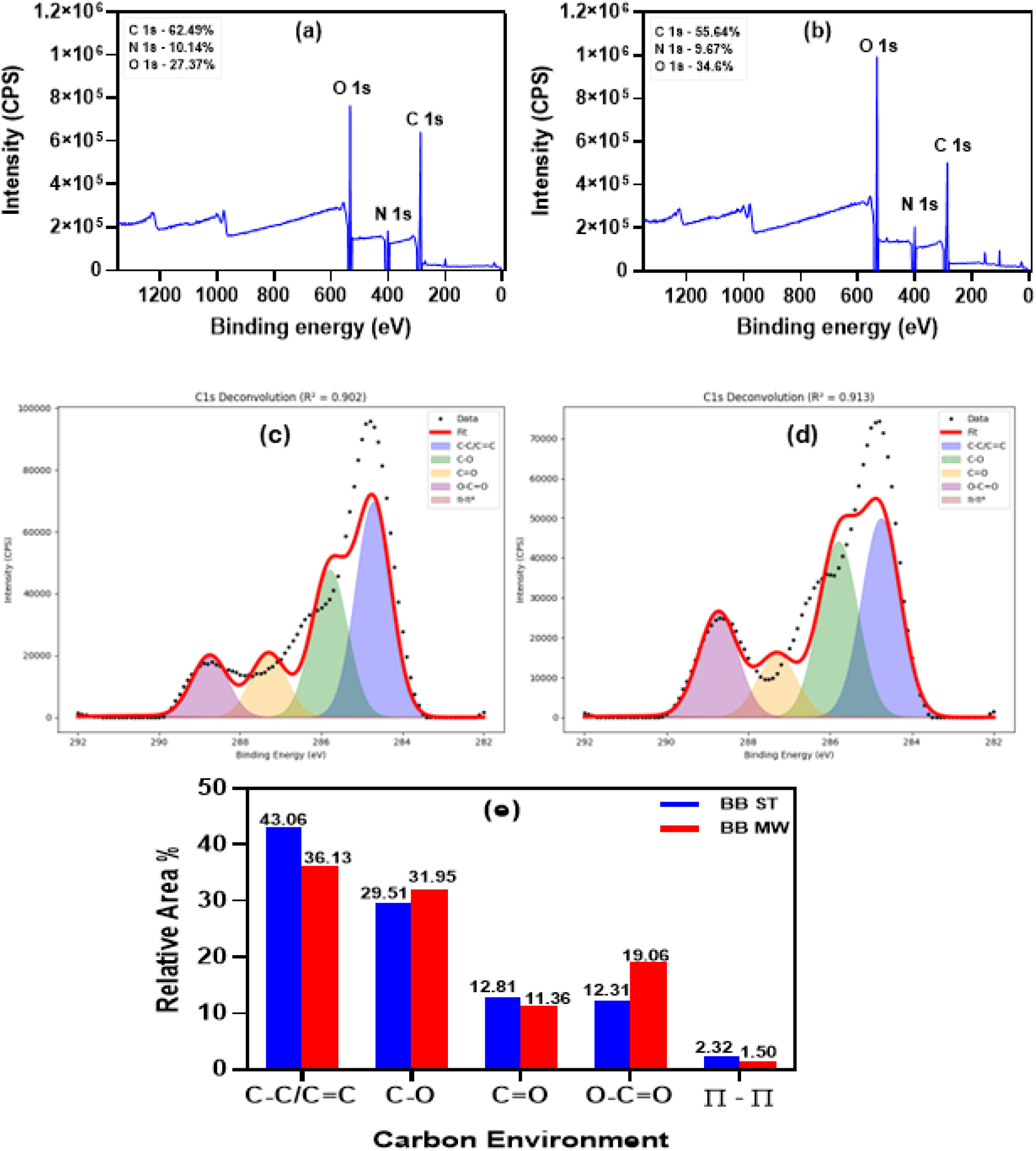
X-ray photoelectron spectroscopic analysis of BB CQDs. XPS survey spectra of (a) BB HT CQDs and (b) BB MW CQDs confirming the presence of C, N, and O; elemental compositions are annotated (BB HT: C 62.49, N 10.14, O 27.37 at.%; BB MW: C 55.64, N 9.67, O 34.60 at.%). High-resolution C 1s spectra with Gaussian deconvolution for (c) BB HT CQDs (R^2^ = 0.902) and (d) BB MW CQDs (R^2^ = 0.913); fitted components: C-C/C=C (284.72/284.75 eV), C-O (285.8 eV), C=O (287.3 eV), O-C=O (288.75/288.73 eV), and π-π* satellite (290.50/290.51 eV). (e) Grouped bar chart comparing the relative area contributions of each carbon environment between BB HT (blue) and BB MW (red) CQDs, highlighting the substantially higher O-C=O fraction and lower graphitic C-C/C=C fraction in the microwave-derived material. See Tables 4 and 5 for full deconvolution parameters.

**Table 3.** XPS elemental composition (at.%) and atomic ratios for BB HT and BB MW CQDs.

| Parameter | BB HT CQDs | BB MW CQDs | Interpretation |
| --- | --- | --- | --- |
| C 1s (at.%) | 62.49 | 55.64 | Higher carbon content in HT reflects greater degree of graphitisation |
| N 1s (at.%) | 10.14 | 9.67 | Comparable nitrogen incorporation from urea in both routes |
| O 1s (at.%) | 27.37 | 34.60 | Higher oxygen in MW attributed to elevated carboxylate surface density |
| C/O atomic ratio | 2.28 | 1.61 | Higher C/O ratio in HT consistent with more graphitised, less oxygenated core |
| C/N atomic ratio | 6.16 | 5.75 | Similar C/N in both routes; nitrogen doping largely route-independent |

The elemental composition of BB-HT CQDs is C 62.49 at.%, N 10.14 at.%, and O 27.37 at.%. This translates to an atomic ratio of 2.28 for C/O. BB-MW CQDs, on the other hand, have a lower C/O ratio of 1.61 since they include C 55.64 at.%, N 9.67 at.%, and O 34.60 at.%. The higher C/O ratio in BB-HT CQDs indicates a greater degree of graphitisation. A larger fraction of carbon is incorporated into aromatic C-C and C=C domains rather than oxygenated functionalities. Nitrogen content remains nearly constant between the two routes (10.14 vs 9.67 at.%). This implies that in the conditions employed, urea-derived nitrogen incorporation is essentially independent of the synthesis process. FTIR data demonstrating similar N-H environments in both samples is consistent with the observation.

The paradox of higher bulk O content in BB MW CQDs (34.60 at.%) despite lower hydroxyl/carbonyl intensity in the FTIR difference spectrum is resolved by the C 1s high-resolution data: the MW product is enriched not in hydroxyl/carbonyl groups but specifically in carboxylate (O-C=O) species, each of which contributes two oxygen atoms to the elemental composition. The XPS C 1s deconvolution results, which are summarized in Tables 4 and 5 and depicted in Figure 6c–e, accurately reflect this.

**Table 4.**
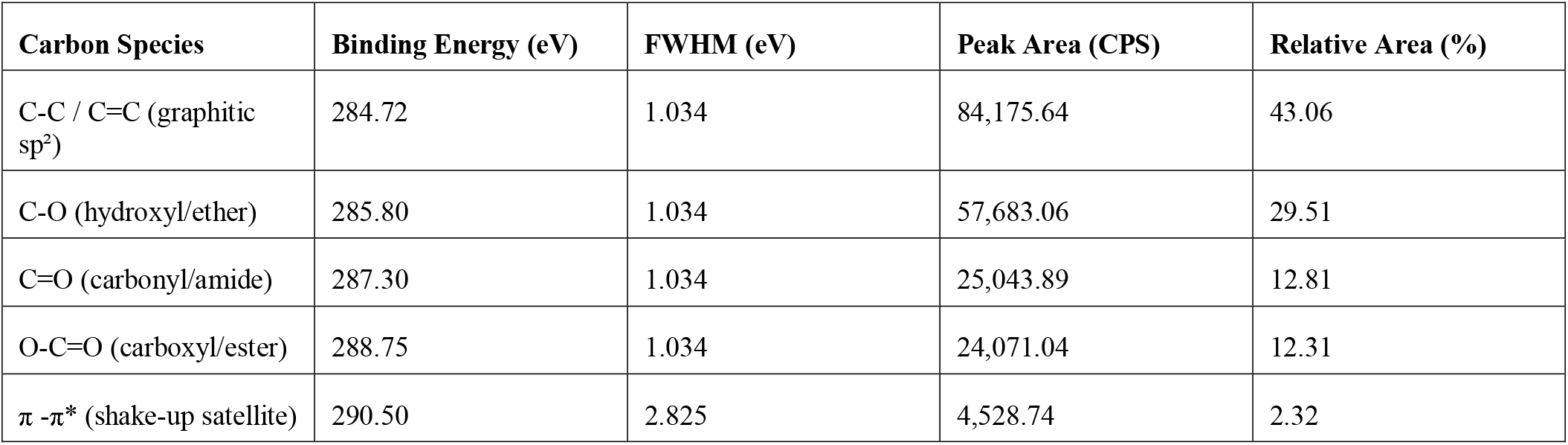
Deconvoluted C 1s XPS peak parameters for BB HT CQDs (five-component Gaussian fit; R^2^ = 0.902).

**Table 5.** Deconvoluted C 1s XPS peak parameters for BB MW CQDs (five-component Gaussian fit; R^2^ = 0.913).

| Carbon Species | Binding Energy (eV) | FWHM (eV) | Peak Area (CPS) | Relative Area (%) |
| --- | --- | --- | --- | --- |
| C-C / C=C (graphitic $sp^2$) | 284.75 | 1.119 | 70,778.02 | 36.13 |
| C-O (hydroxyl/ether) | 285.80 | 1.119 | 62,600.94 | 31.95 |
| C=O (carbonyl/amide) | 287.30 | 1.119 | 22,251.23 | 11.36 |
| O-C=O (carboxyl/ester) | 288.73 | 1.119 | 37,349.20 | 19.06 |
| $\pi - \pi^*$ (shake-up satellite) | 290.51 | 2.825 | 2,934.15 | 1.50 |

For BB HT CQDs, the dominant C 1s component is C-C/C=C at 284.72 eV (43.06%), confirming the presence of a graphitic sp^2^ carbon core. Hydroxylic C-O at 285.8 eV accounts for 29.51%, whilst carbonyl C=O (287.3 eV, 12.81%), carboxyl/ester O-C=O (288.75 eV, 12.31%), and π-π* shake-up satellite (290.50 eV, 2.32%) together constitute the oxygenated minority. The graphitic C-C/C=C proportion in BB MW CQDs is significantly lower (36.13%), however the O-C=O component rises dramatically to 19.06%, a 6.75 percentage-point increase that is the most diagnostically significant difference between the two materials. Additionally, the MW product has a slightly larger C-O percentage (31.95%).

The underlying mechanism is straightforward. Under microwave conditions, decarboxylation of the citric acid-derived precursor proceeds incompletely, leaving a larger proportion of carboxylate groups intact at the particle surface. Prolonged hydrothermal treatment drives this process further towards completion. Carboxylate species are progressively converted into carbonyl C=O environments and, ultimately, into graphitic C–C/C=C domains as the reaction advances. The reduced π-π* satellite in MW CQDs (1.50% vs. 2.32%) independently corroborates the lower degree of aromatic conjugation in the microwave product. The FTIR difference-spectroscopy data stated above are quantitatively supported by this XPS evidence, which is entirely consistent with it.

### 3.5 In Vitro Cytotoxicity

MTT assays were performed in RPE-1 and HeLa cells across 10-250 µg/mL after 24 h exposure (Figure 7). Both CQD preparations were non-cytotoxic across the entire concentration range tested in both cell lines, with no statistically significant reduction in viability relative to untreated controls at any dose^23^.

**Figure 7.**
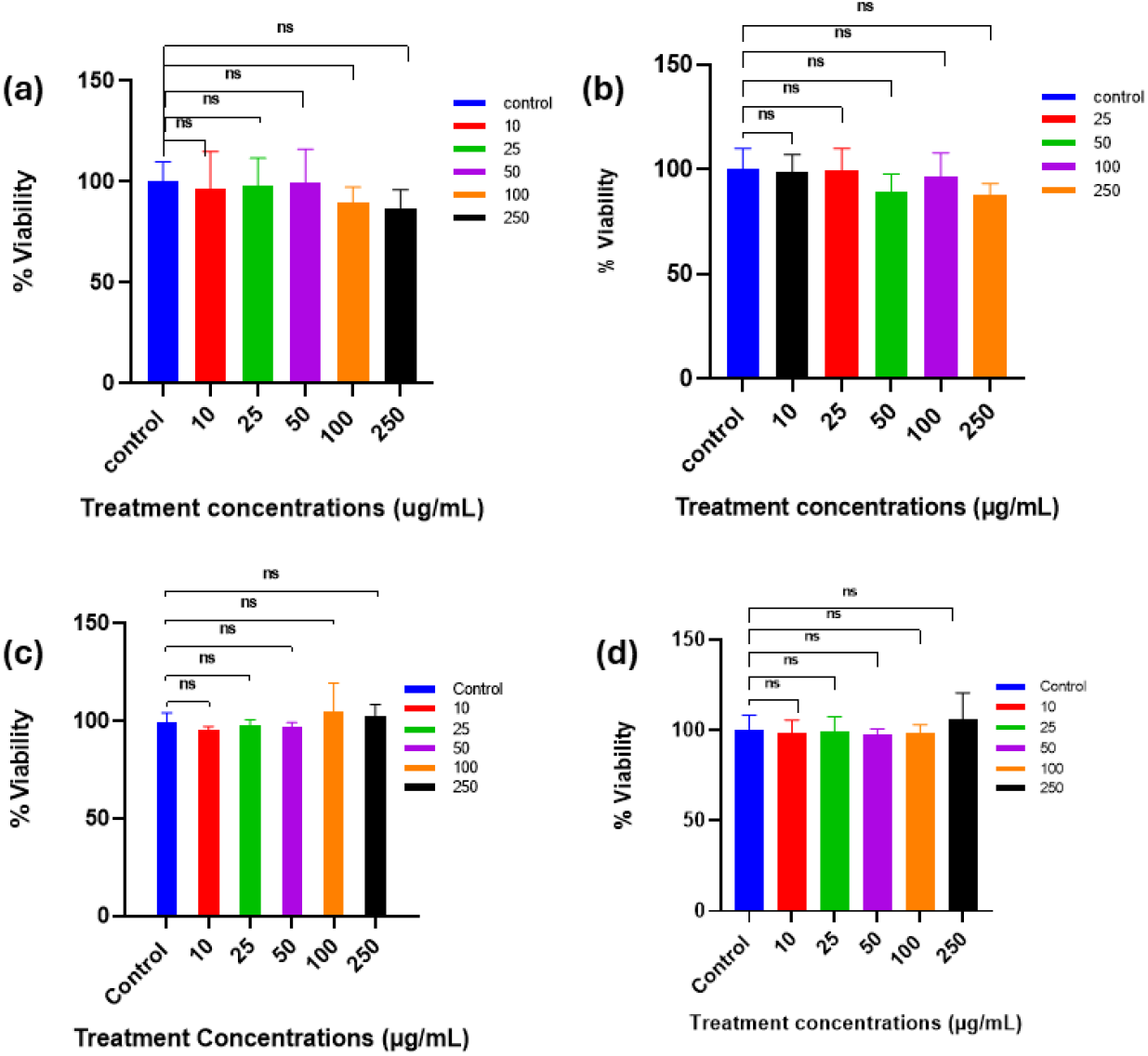
In vitro cytotoxicity of BB CQDs assessed by MTT assay after 24 h exposure (10,000 cells/well, 96-well format). Percentage viability relative to untreated control for (a) BB HT CQDs in RPE-1 cells (10–250 µg/mL), (b) BB MW CQDs in RPE-1 cells (25–250 µg/mL), (c) BB HT CQDs in HeLa cells (10–250 µg/mL), and (d) BB MW CQDs in HeLa cells (10–250 µg/mL). Data are expressed as mean ± SD (n = 3). Statistical significance compared with the untreated control was evaluated using one-way ANOVA followed by Dunnett’s post hoc test: ns, p > 0.05. No statistically significant cytotoxicity was observed at any tested concentration in either cell line for either preparation.

In RPE-1 cells, BB HT CQDs maintained viability between 90-99% across all concentrations (Figure 7a). The modest numerical decline at 100 µg/mL (~90%) and 250 µg/mL (~87%) did not reach statistical significance (p > 0.05). BB MW CQDs showed a comparable profile in RPE-1 cells, with viability ranging from ~85–100% across 25–250 µg/mL, again entirely non-significant (Figure 7b). Minor numerical variations between groups reflect biological scatter rather than dose-dependent toxicity.

In HeLa cells, both preparations produced viability values that remained within or above the control range throughout. BB HT CQDs fluctuated between ~95–103% of control across 10–250 µg/mL (Figure 7c). BB MW CQDs showed a similarly flat response, with values spanning ~97–105% (Figure 7d). The slight apparent elevation above 100% at certain concentrations is a commonly observed phenomenon in MTT assays attributable to CQD-associated metabolic stimulation or mild formazan interference at sub-cytotoxic doses it does not indicate cytoproliferation and carries no biological significance here.

The consistent non-significance across both cell lines and both preparations establishes a clean biocompatibility profile up to 250 µg/mL. This concentration substantially exceeds the doses at which both functional properties operate: the antioxidant IC□□ of BB HT CQDs is 38.48 µg/mL, and statistically significant zebrafish uptake of BB MW CQDs begins at 100 µg/mL. There is therefore a clear safety margin of at least two-to sixfold between the effective functional dose and the highest tested concentration, which produced no measurable harm to either cell type.

The absence of differential toxicity between RPE-1 and HeLa cells despite the markedly different uptake profiles documented in Section 3.10 is mechanistically consistent. Greater CQD internalisation in HeLa cells does not translate into greater cytotoxicity because the transformed cell line maintains elevated antioxidant buffering capacity through upregulated glutathione and thioredoxin systems^24^. These systems can accommodate the redox load imposed by internalised CQDs at the concentrations used here. RPE-1 cells, which take up fewer particles, are correspondingly exposed to lower intracellular particle burdens further reason why neither cell type reaches a cytotoxic threshold within the tested range^25^.

### 3.6 DPPH Radical Scavenging Activity

Dose-dependent DPPH radical scavenging was measured across 25–1000 µg/mL for both CQD preparations and benchmarked against ascorbic acid as a positive reference standard (**Figure 8**). All three materials showed progressive, concentration-dependent inhibition that approached saturation at 1000 µg/mL.

**Figure 8.**
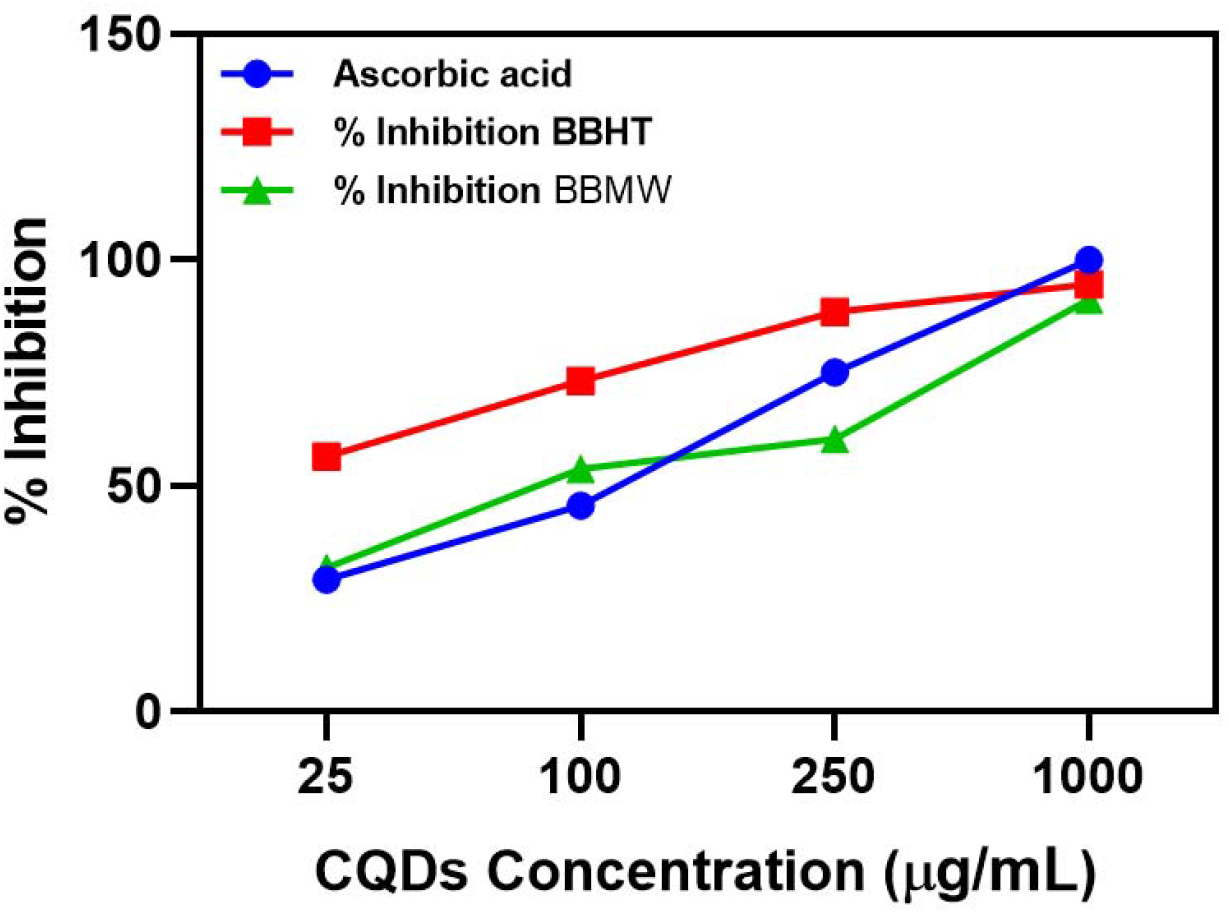
DPPH radical scavenging activity of BB CQDs benchmarked against ascorbic acid. Percentage DPPH inhibition for BB HT CQDs (red), BB MW CQDs (green), and ascorbic acid positive control (blue) at concentrations of 25, 100, 250, and 1000 µg/mL. BB HT CQDs exceed ascorbic acid inhibition at 25, 100, and 250 µg/mL, converging at 1000 µg/mL (~97% vs ~99%). IC□□values determined by nonlinear four-parameter regression: BB HT CQDs = 38.48 µg/mL; BB MW CQDs = 109.9 µg/mL. Data represent mean ± SD (n = 3).

BB HT CQDs were the most potent radical scavengers across the tested range. At 25 µg/mL, they achieved ~57% inhibition exceeding both ascorbic acid (~32%) and BB MW CQDs (~38%) at the same concentration. This hierarchy was maintained at 100 µg/mL (BB HT ~72%, ascorbic acid ~55%, BB MW ~52%) and 250 µg/mL (BB HT ~87%, ascorbic acid ~75%, BB MW ~60%). All three converged at 1000 µg/mL, reaching ~92–99% inhibition. The calculated IC□□ values are 38.48 µg/mL for BB HT CQDs and 109.9 µg/mL for BB MW CQDs 2.86-fold difference in potency. That BB HT CQDs outperform ascorbic acid below 1000 µg/mL is a result of direct mechanistic relevance, given that ascorbic acid is a clinically used antioxidant standard.

The antioxidant superiority of BB HT CQDs is grounded in two structural features identified by XPS and FTIR. The elevated surface density of hydroxyl and amine groups (FTIR, 3438 cm□^1^) supports hydrogen-atom transfer (HAT), in which O–H and N–H groups donate hydrogen atoms to DPPH radicals, producing resonance-stabilised radicals delocalised across the graphitic core^26^. The higher graphitic C–C/C=C fraction (XPS, 43.06%) enables a parallel single-electron transfer (SET) pathway, where the extended π-electron network donates electrons directly to DPPH^22^. These two mechanisms act simultaneously, which likely explains why BB HT CQDs reach ~57% inhibition at just 25 µg/mL concentration at which ascorbic acid achieves only ~32%.

BB MW CQDs, despite their lower potency, are not without antioxidant activity. At 1000 µg/mL they reach ~92% inhibition, close to the ascorbic acid ceiling. Their lower carboxylate-dominated surface provides fewer HAT-active proton donors, which suppresses activity at sub-100 µg/mL concentrations. The crossover point where BB MW CQDs match ascorbic acid occurs between 100 and 1000 µg/mL, consistent with their higher IC□□.

The partial incorporation of bearberry phytochemicals particularly arbutin-derived hydroquinone and galloyl-type structures is likely enhanced under sustained hydrothermal conditions relative to the brief microwave process. These species contribute additional phenolic O–H donors that augment the intrinsic radical-scavenging capacity of the CQD surface, and their preferential retention in BB HT CQDs is consistent with the known thermal stability of gallotannins and ellagitannins at 150–180 °C^27^.

### 3.7 In Vivo Fluorescence Uptake in Zebrafish Larvae

For zebrafish imaging, BB MW CQDs were chosen because of their higher quantum yield, which offers a larger signal margin over larval autofluorescence. Larvae at 72 hpf were exposed to 50-500 µg/mL for 24 h and imaged across brightfield, fluorescence, and merged channels (Figure 9). Brightfield images show structurally normal larvae at every concentration. Somite patterning, notochord integrity, and head morphology are preserved throughout, with no curvature defects, pericardial oedema, or yolk sac abnormalities. CQD exposure at these doses does not impair larval development over the 4 h window^7,28^. The fluorescence channel tells a different story at each concentration. The control and 50 µg/mL groups are visually indistinguishable only baseline autofluorescence is present. At 100 µg/mL, green emission is clearly elevated across the trunk and abdominal compartment. By 250 µg/mL, fluorescence is distributed broadly through the larval body, with particularly intense signal localised to the yolk sac and intestinal cavity. At 500 µg/mL, this signal approaches channel saturation in the abdominal region. Merged pictures rule out simple particle surface adhesion by confirming tissue co-localization at all positive doses.

**Figure 9.**
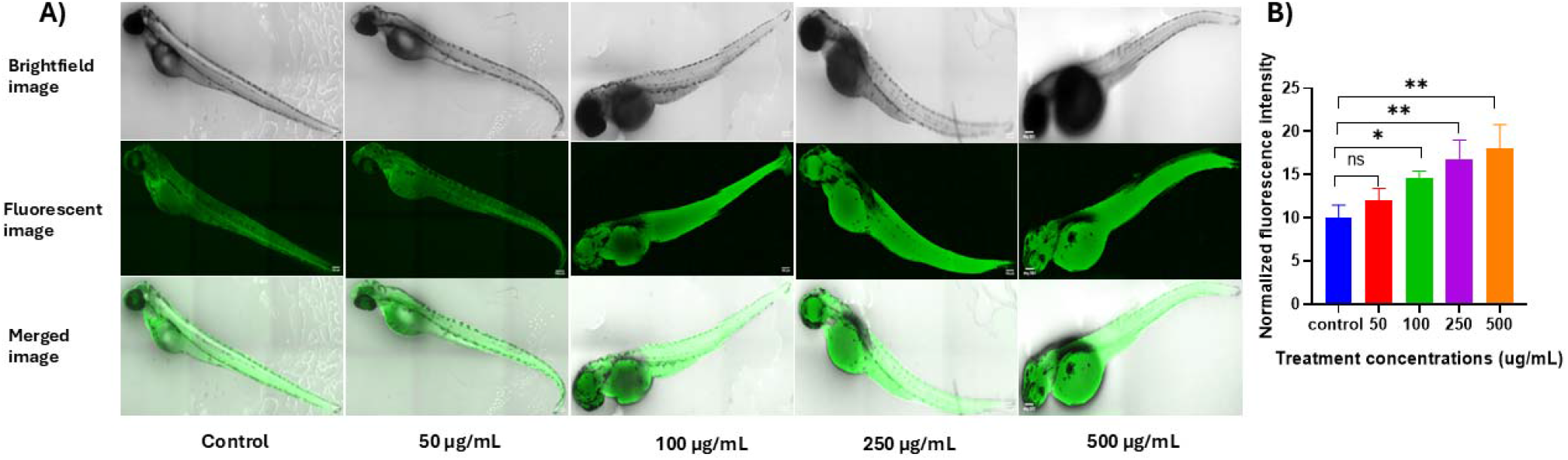
In vivo fluorescence uptake of BB MW CQDs in zebrafish larvae. (A) Representative brightfield (top), fluorescence (middle), and merged composite (bottom) images of 72 hpf larvae after 4 h exposure to CQDs at 50, 100, 250, and 500 µg/mL BB MW CQDs. Brightfield images confirm morphological integrity across all treatment groups. Fluorescence images reveal a progressive, concentration-dependent increase in green channel emission, with signal distributed across the larval trunk and concentrated in the yolk sac and intestinal compartment at 250 and 500 µg/mL. Scale bars: 500 µm. (B) Normalised whole-larval fluorescence intensity quantified in ImageJ from background-corrected regions of interest. Results are presented as mean ± SD (n = 3 per group). Statistical differences relative to the untreated control were determined using one-way ANOVA followed by Dunnett’s post hoc multiple-comparison test: ns, p > 0.05; *, p < 0.05; **, p < 0.01. Significant accumulation is confirmed from 100 µg/mL (p < 0.05), with plateau behaviour and increased inter-individual variability evident at 500 µg/mL.

Quantitative analysis supports the visual pattern. Normalised fluorescence rises from ~10 a.u. at control to ~11 a.u. at 50 µg/mL (ns), ~13–14 a.u. at 100 µg/mL (p < 0.05), ~16.5 a.u. at 250 µg/mL (p < 0.01), and ~17.5–18 a.u. at 500 µg/mL (p < 0.01). The plateau between 250 and 500 µg/mL, alongside widening error bars at the highest dose, suggests that uptake approaches saturation and that inter-individual variation in gill and integument permeability becomes the dominant source of variability at high particle burdens. The non-significance at 50 µg/mL is most plausibly a sensitivity limit of whole-larval mean grey value quantification rather than evidence of zero internalisation. Confocal or light-sheet imaging of individual organs at this concentration would likely resolve CQD-specific signal clearly. Three physicochemical properties drive the observed uptake. The 9.65 nm hydrodynamic diameter of BB MW CQDs, confirmed by DLS, falls within the size range permitting passive diffusion across the larval integument and intestinal epithelium without requiring active transport. The PEG 2000 surface layer suppresses opsonisation, extending circulatory residence time and enabling systemic tissue distribution rather than rapid immune sequestration. The high quantum yield of BB MW CQDs enabled strong fluorescence detection even at relatively low tissue concentrations, exceeding the intrinsic autofluorescence background of zebrafish larvae^29^. This property partly motivated the selection of BB MW CQDs over BB HT CQDs for in vivo imaging studies. Accumulation within the yolk sac at 250 and 500 µg/mL agrees with reported uptake patterns of hydrophilic sub-10 nm nanoparticles in zebrafish larvae, where the yolk syncytial layer functions as a major site of nanoparticle deposition following intestinal absorption^30^. At all investigated concentrations, no morphological harm was seen. These results justify additional preclinical testing and extend the favorable in vitro biocompatibility profile into a real vertebrate model.

### 3.8 Dynamic Light Scattering and Zeta Potential

Both CQD samples display narrow, monomodal patterns within 10 nm, according to hydrodynamic size distributions derived by DLS (Figure 10). This is consistent with the expected size range for carbon quantum dots synthesised via hydrothermal and microwave methods. BB-HT CQDs display an intensity-weighted mean hydrodynamic diameter of 7.13 nm. In comparison, BB-MW CQDs show a slightly larger mean diameter of 9.65 nm. The larger hydrodynamic size of the microwave-derived particles can be attributed to their higher surface carboxylate density. XPS C 1s analysis confirms a greater O-C=O contribution in BB-MW CQDs (19.06%) relative to BB-HT CQDs (12.31%). Carboxylate-rich surfaces tend to retain a more extended hydration shell, which increases the apparent hydrodynamic radius beyond the physical core size. Additionally, structural variations in the carbon core are consistent with this observation. The graphitic C-C/C=C proportion is smaller in BB-MW CQDs (36.13%) than in BB-HT CQDs (43.06%). This implies that the microwave-derived material has a less compact and less graphitized core.

**Figure 10.**
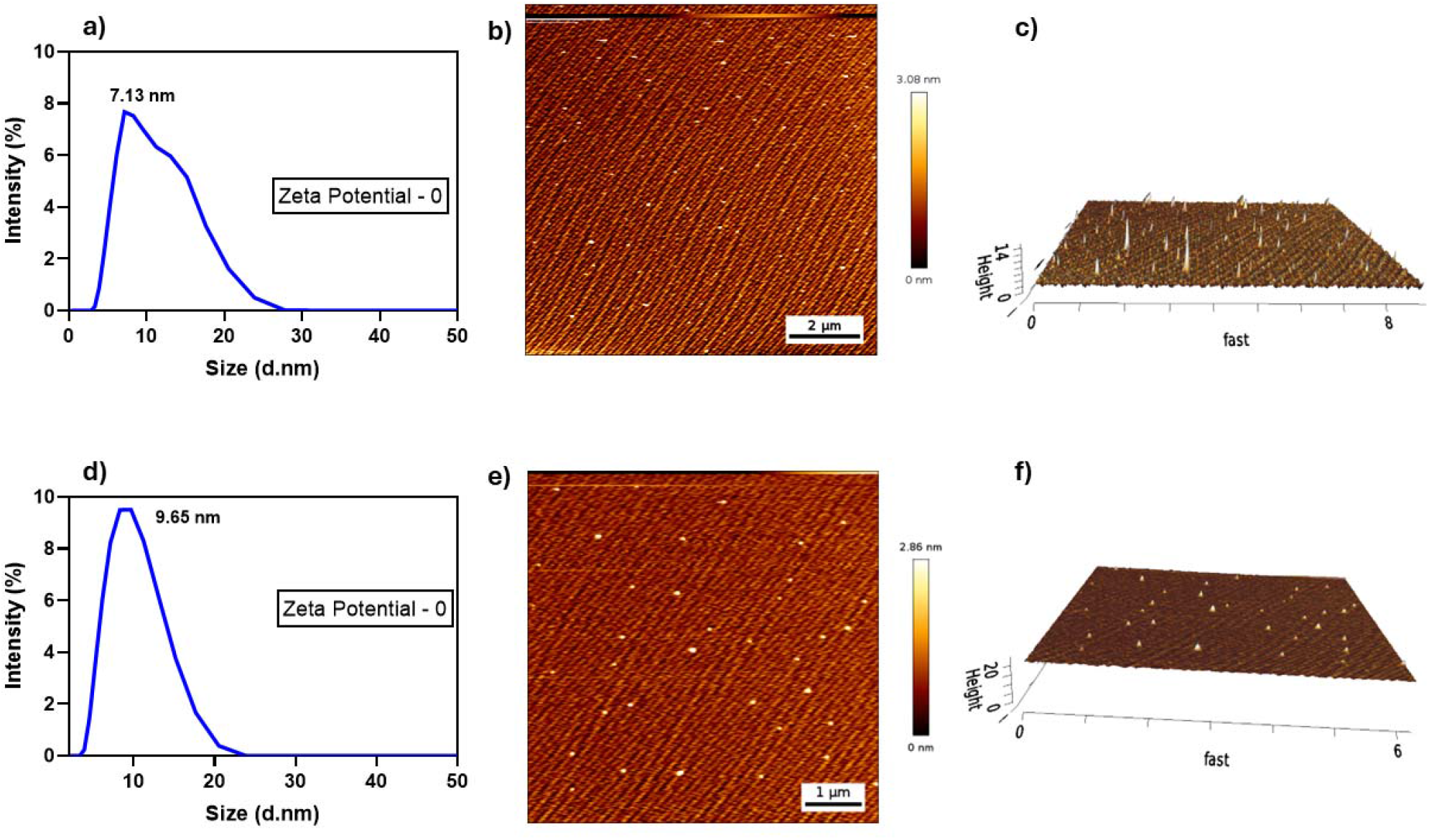
Dynamic light scattering and atomic force microscopy characterisation of BB CQDs. Intensity-weighted hydrodynamic size distributions for (a) BB HT CQDs (mean diameter 7.13 nm) and (d) BB MW CQDs (mean diameter 9.65 nm) measured in deionised water at 25 °C; both preparations display narrow, monomodal profiles indicative of well-dispersed, non-aggregated colloidal suspensions. Near-zero zeta potential values (~0 mV) confirm steric rather than electrostatic colloidal stabilisation by the PEG 2000 surface layer. Two-dimensional AFM topography height maps of (b) BB HT CQDs (scan area 10 × 10 µm; scale bar 2 µm; height scale 0–3.08 nm) and (e) BB MW CQDs (scan area 5 × 5 µm; scale bar 1 µm; height scale 0–2.86 nm) reveal discrete, quasi-spherical nanoparticles with no evidence of aggregation or anisotropic morphologies. Three-dimensional topographic renders of (c) BB HT CQDs and (f) BB MW CQDs further confirm the dot-like particle geometry and the greater surface density and marginally higher topographic height of the hydrothermally derived material. The larger proportional reduction between DLS diameter and AFM height in BB MW CQDs (70%) relative to BB HT CQDs (57%) is attributed to the lower degree of core graphitisation and greater elastic compressibility of the microwave-derived material, consistent with XPS C 1s deconvolution data.

Zeta potential measurements yielded near-zero values (≈0 mV) for both BB HT and BB MW CQDs. The dense PEG 2000 corona on both CQD surfaces creates an entropic repulsion barrier that prevents inter-particle coalescence independently of surface charge^31^. This is confirmed by the narrow monomodal DLS profiles observed in both preparations. The low zeta potential further implies that these materials will be relatively insensitive to ionic-strength-dependent aggregation in physiological media a valuable property for biological applications in cell culture medium and in vivo environments where high electrolyte concentrations would destabilise electrostatically stabilised nanoparticles.

The combined DLS and XPS data provide a coherent structural interpretation. BB-HT CQDs exhibit a smaller, more graphitised core with lower surface carboxylate density. This corresponds to a reduced hydrodynamic diameter. In contrast, BB-MW CQDs display a less condensed carbon core and higher carboxylate coverage. This results in a modestly increased hydrodynamic size due to an expanded hydration shell. Steric stabilisation from the PEG 2000 surface layer prevents aggregation and ensures good dispersion in aqueous media^32^.

### 3.9 Atomic Force Microscopy

AFM topography images and three-dimensional renders for BB HT and BB MW CQDs are presented in **Figure 10**. Both preparations display the canonical morphological signature of carbon quantum dots: discrete, quasi-spherical protrusions of sub-5 nm height deposited on an atomically flat mica substrate, with no evidence of rod-like, sheet-like, or significantly elongated nanostructures. The absence of large aggregates in either preparation corroborates the narrow, monomodal DLS distributions and confirms that the steric stabilisation conferred by the PEG 2000 surface layer remains effective upon deposition from dilute aqueous suspension.

The maximum topographic heights extracted from the AFM colour scales are 3.08 nm for BB HT CQDs and 2.86 nm for BB MW CQDs. Two physical factors collectively account for the DLS-AFM size discrepancy in both preparations. First, DLS reports an intensity-weighted hydrodynamic diameter that includes the fully hydrated PEG 2000 corona and its associated bound water shell contributions that are entirely absent from the dry, vacuum-deposited state sampled by AFM. Second, upon drying onto the mica substrate, the flexible PEG chains collapse and flatten radially, reducing the effective height contribution of the surface layer from the hydrodynamic measurement^**33**^.

The larger proportional DLS-AFM discrepancy observed for BB MW CQDs (70% reduction) relative to BB HT CQDs (57% reduction) is mechanistically coherent when interpreted against the XPS compositional data. BB HT CQDs possess a more graphitised carbon core (C–C/C=C fraction: 43.06%) characterised by densely stacked sp^2^-hybridised aromatic domains, which confer greater mechanical rigidity and resistance to tip-induced compression^**22**^. BB MW CQDs, with their lower graphitic fraction (36.13%) and correspondingly higher proportion of sp^3^-hybridised and carboxylate-bearing carbon environments, possess a comparatively softer, less mechanically consolidated core that undergoes greater elastic deformation under the AFM tip^**34**^. Furthermore, it is anticipated that the higher carboxylate surface density of BB MW CQDs (O–C=O: 19.06% vs. 12.31% for HT) will support a longer hydration shell in solution in accordance with their larger DLS diameter. However, this longer shell dehydrates and collapses more completely upon drying, which contributes disproportionately to the height reduction seen between wet (DLS) and dry (AFM) measurements. When considered collectively, the higher core compactness and surface rigidity of the hydrothermally generated material are directly expressed topographically by the AFM height ratio of BB HT > BB MW, despite the reverse ordering in DLS diameter.

Two morphological differences between the preparations are also evident in the 2D topography images. BB HT CQDs exhibit a markedly higher particle surface density per unit scan area, with numerous closely spaced but individually resolved protrusions visible across the 10 × 10 µm field. BB MW CQDs, imaged over a smaller 5 × 5 µm scan area, show a sparser, more isolated particle distribution. At equivalent deposited mass concentration, a higher number density of BB HT CQDs per unit area is consistent with their smaller physical size, which means more particles are present per unit mass. The three-dimensional renders reinforce these observations: BB HT CQDs appear as a dense field of sharp, spike-like protrusions of relatively uniform height, whilst BB MW CQDs form a sparser landscape of slightly lower, more rounded protrusions, the latter morphology again consistent with the softer, less graphitised core undergoing a slightly greater degree of deformation during deposition and drying.

In summary, AFM analysis confirms the sub-5 nm physical size, quasi-spherical morphology, and non-aggregated dispersion state of both CQD preparations, whilst the route-dependent difference in core graphitisation identified by XPS manifests directly in the topographic height and DLS-AFM compression ratio of the two materials. These observations complete a self-consistent, multi-technique physical characterisation in which DLS, XPS, FTIR, and AFM all independently converge on the same mechanistic narrative: hydrothermal synthesis produces a smaller, harder, more graphitised particle with higher surface hydroxyl and carbonyl density, whilst microwave synthesis yields a slightly larger, softer particle with a carboxylate-dominated surface differences that propagate coherently into all measured optical, antioxidant, and biological properties.

### 3.10 Cellular Uptake in Non-Cancerous and Cancer Cell Lines

Confocal fluorescence microscopy was used to examine intracellular CQD uptake in RPE-1 and HeLa cells after 24 h exposure to 100, 200, and 300 µg/mL CQDs (**Figures 11–14**). Fluorescence intensities normalised to untreated controls are summarised in **Figure 15**. Clear differences in uptake behaviour were observed between the two cell lines. HeLa cells consistently showed stronger intracellular fluorescence than RPE-1 cells at all tested concentrations. In both models, BB MW CQDs also produced higher fluorescence signals than BB HT CQDs.

**Figure 11.**
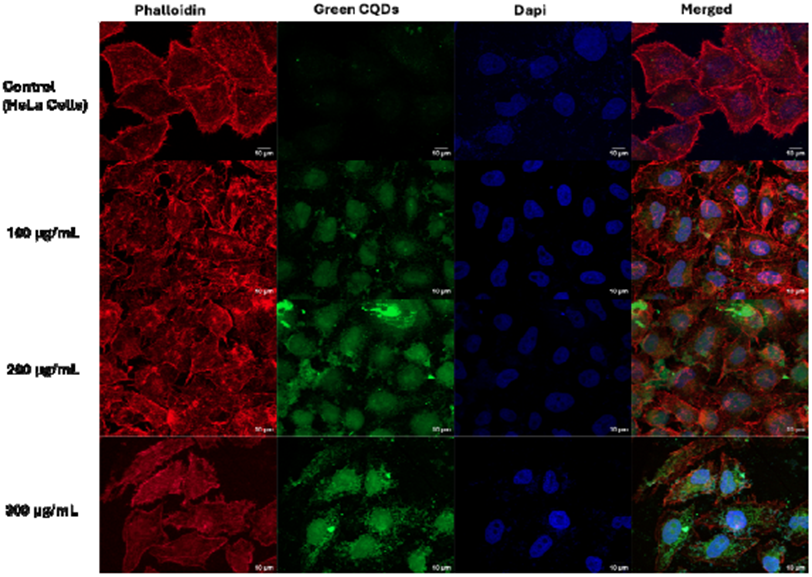
Confocal fluorescence microscopy of BB HT CQD uptake in HeLa cervical carcinoma cells. Images show (columns, left to right) phalloidin–FITC staining of F-actin (green), CQD emission channel (cyan/green), DAPI nuclear counterstain (blue), and merged composite at (rows, top to bottom) untreated control, 100, 200, and 300 µg/mL BB HT CQDs after 24 h exposure. Scale bars: 20 µm. CQD fluorescence colocalises with intracellular compartments, indicating genuine cellular internalisation. Concentration-dependent increase in intracellular CQD signal is evident in the merged panels; all concentrations show marginal statistical trend (p < 0.1.

**Figure 12.**
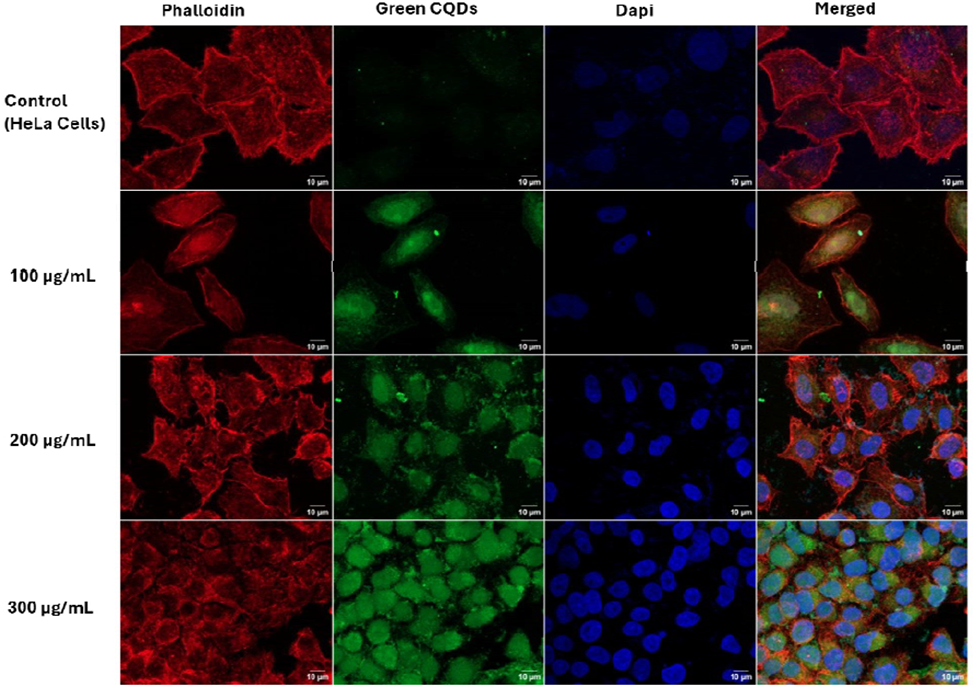
Confocal fluorescence microscopy of BB MW CQD uptake in HeLa cervical carcinoma cells. Imaging layout identical to Figure 11: phalloidin–FITC (actin, green), CQD emission (cyan/green), DAPI (nuclei, blue), and m rged composite at untreated control, 100, 200, and 300 µg/mL BB MW CQDs after 24 h. Scale bars: 20 µm. BB MW CQDs produce greater absolute intracellular fluorescence than BB HT CQDs across all concentrations, reaching approximately 700% of control at 300 µg/mL, consistent with the higher quantum yield and surface carboxylate density of the microwave-derived material. All concentrations show a marginal upward trend (p < 0.1) relative to control.

**Figure 13.**
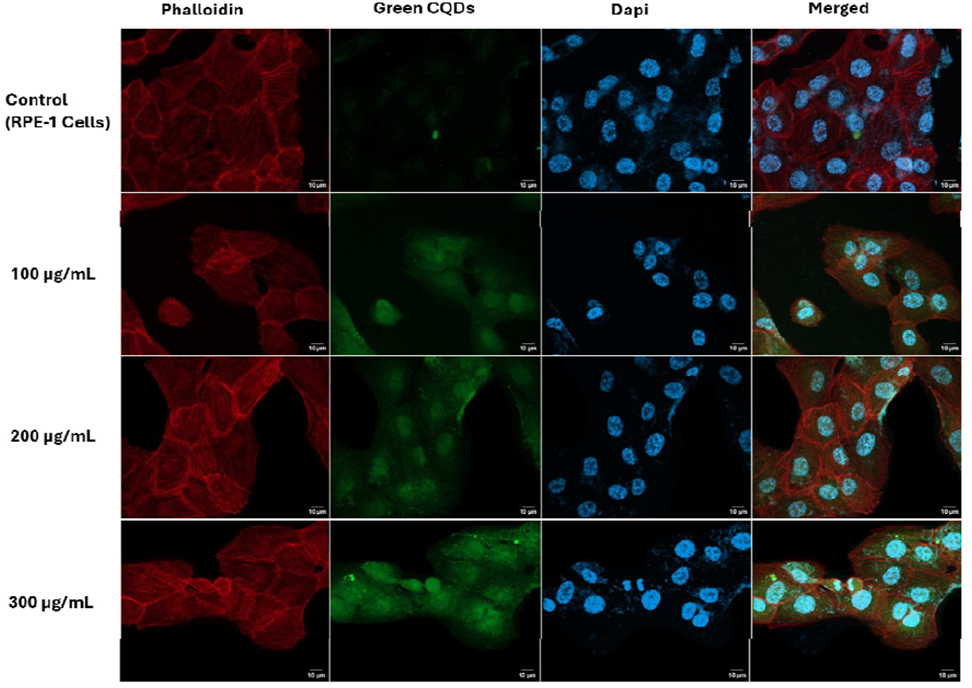
Confocal fluorescence microscopy of BB HT CQD uptake in RPE-1 non-cancerous retinal pigment epithelial cells. Images show phalloidin–FITC (actin, green), CQD emission (cyan/green), DAPI (nuclei, blue), and merged composite at untreated control, 100, 200, and 300 µg/mL BB HT CQDs after 24 h exposure. Scale bars: 20 µm. Intracellular CQD fluorescence does not reach statistical significance at 100 or 200 µg/mL (p > 0.05) and shows only a marginal trend at 300 µg/mL (p < 0.1), consistent with the lower endocytic activity of this non-cancerous cell line.

**Figure 14.**
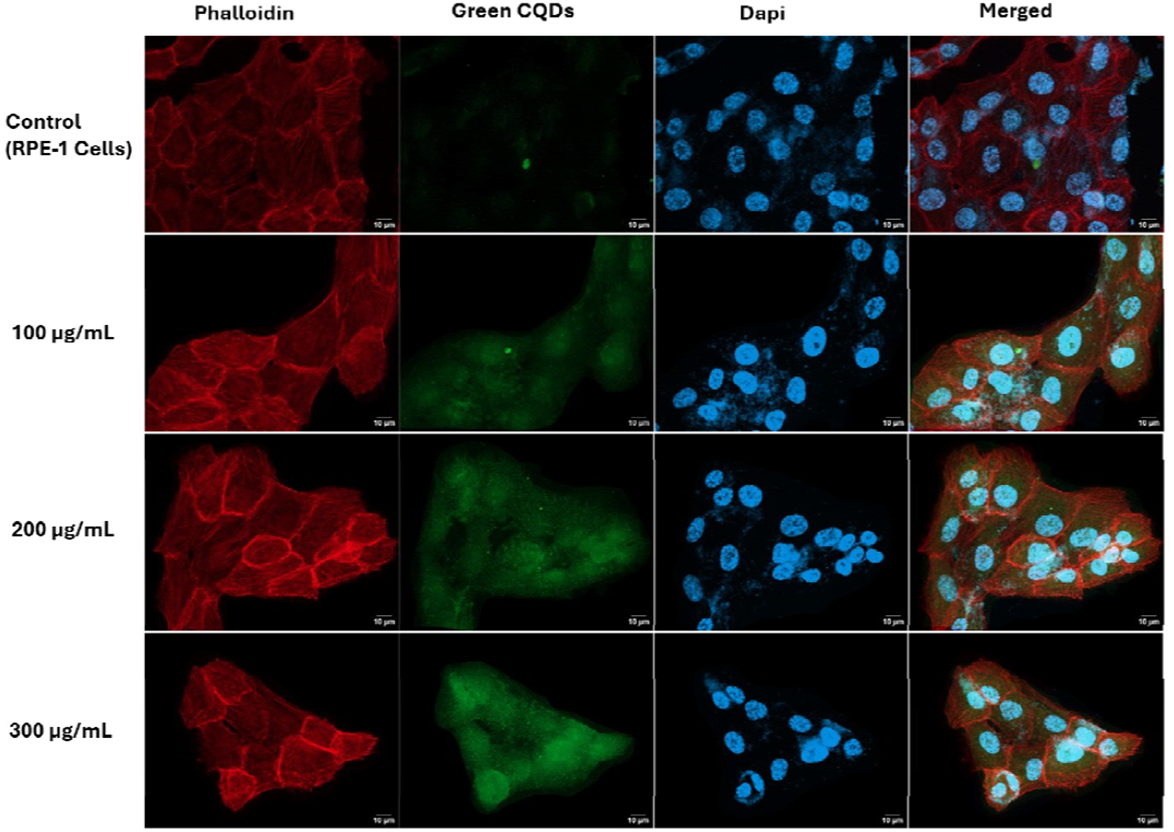
Confocal fluorescence microscopy of BB MW CQD uptake in RPE-1 non-cancerous retinal pigment epithelial cells. Imaging layout as per Figures 11–13. BB MW CQDs produce a statistically significant (p < 0.001) increase in intracellular fluorescence at 200 µg/mL in RPE-1 cells, as indicated by the *** annotation. A marginal upward trend is observed at 300 µg/mL (p < 0.1, #). The high-significance uptake at 200 µg/mL is attributed to the elevated surface carboxylate density of BB MW CQDs (O–C=O: 19.06%), which promotes receptor-mediated and endocytic internalisation pathways. Scale bars: 20 µm.

**Figure 15.**
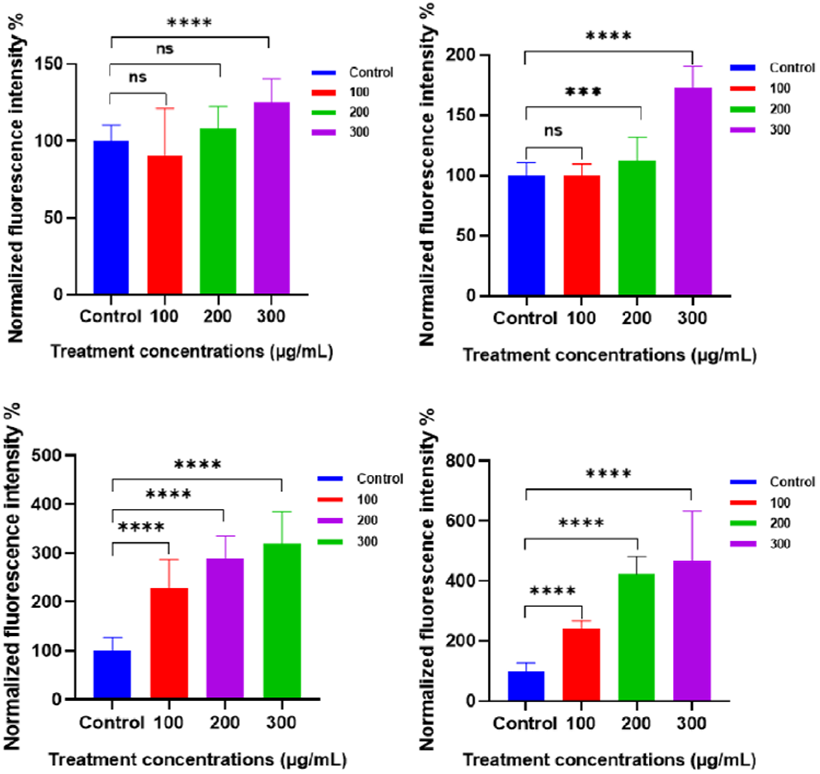
Quantitative analysis of intracellular CQD uptake by confocal fluorescence microscopy. Normalised fluorescence intensity (% of untreated control) in RPE-1 cells treated with (top-left) BB HT CQDs and (top-right) BB MW CQDs, and in HeLa cells treated with (bottom-left) BB HT CQDs and (bottom-right) BB MW CQDs, at concentrations of 100, 200, and 300 µg/mL after 24 h exposure. Note the different y-axis scales across panels (150%, 200%, 500%, and 800% respectively), reflecting the markedly greater uptake efficiency in HeLa relative to RPE-1 cells. Data represent mean ± SD (n = 3). Statistical comparisons against the untreated control group were performed using one-way ANOVA followed by Dunnett’s multiple-comparison test: ns, p > 0.05; ***, p < 0.001; ****, p < 0.0001. Both CQD preparations achieve statistically significant (p < 0.0001) intracellular accumulation in HeLa cells from 100 µg/mL, whilst RPE-1 cells require higher concentration thresholds (≥ 200 µg/mL for BB MW CQDs; 300 µg/mL for BB HT CQDs) to reach significance. BB MW CQDs consistently produce greater intracellular fluorescence than BB HT CQDs in both cell lines, reaching approximately 4.7-fold above control in HeLa cells at 300 µg/mL.

In non-cancerous RPE-1 cells, uptake of BB HT CQDs was limited and strongly concentration-dependent. No significant increase in intracellular fluorescence was observed at 100 or 200 µg/mL (p > 0.05), with signal intensity remaining within 10% of the control. A significant increase was detected only at 300 µg/mL (~125% of control; p < 0.0001) indicating a relatively high threshold for CQD internalisation in this non-transformed cell line. BB MW CQDs showed a more efficient and progressive uptake profile in RPE-1 cells. Fluorescence remained comparable to control levels at 100 µg/mL (p > 0.05), but increased significantly at 200 µg/mL (~112%; p < 0.001) and further at 300 µg/mL (~170%; p < 0.0001). The lower uptake threshold of BB MW CQDs in RPE-1 cells, relative to BB HT CQDs, is consistent with their higher surface carboxylate content (O–C=O: 19.06% vs. 12.31% based on XPS C 1s deconvolution). Elevated surface carboxylation is known to facilitate receptor-mediated endocytic uptake.

In HeLa cells the uptake dynamics were fundamentally different. Both CQDs formulations produced significant intracellular fluorescence accumulation from the lowest tested concentration of 100 µg/mL (p < 0.0001). Fluorescence intensity increased progressively at 200 and 300 µg/mL. BB HT CQDs reached approximately 230%, 290%, and 320% of control fluorescence at 100, 200, and 300 µg/mL respectively. BB MW CQDs produced even greater accumulation, reaching approximately 245%, 420%, and 470% of control at the same concentrations. The notably steeper concentration response relationship of BB MW CQDs in HeLa cells with fluorescence nearly doubling between 100 and 200 µg/mL distinguishes this preparation as the more efficient imaging agent in the cancer cell context.

The cell-line selectivity is quantitatively striking. At 300 µg/mL, intracellular fluorescence in HeLa cells was approximately 2.6-fold higher than in RPE-1 cells for BB HT CQDs and 2.8-fold higher for BB MW CQDs. This difference is consistent with the enhanced endocytic activity of cancer cells including elevated clathrin-mediated endocytosis and macropinocytosis which support the high metabolic demands of rapidly proliferating tumours^35^. In contrast non-cancerous RPE-1 cells maintain more tightly regulated endocytic pathways, limiting nanoparticle uptake at lower concentrations. The significant but comparatively restricted uptake observed in RPE-1 cells at 300 µg/mL further confirms the acceptable biocompatibility window established by MTT data (Section 3.5), since functionally relevant imaging concentrations will be substantially lower than those producing measurable cytotoxicity^36,37^.

The consistently superior cellular uptake of BB MW CQDs over BB HT CQDs in both cell lines is mechanistically coherent with two independent physicochemical factors. First, the higher carboxylate surface coverage of the microwave-derived material engages membrane-associated transport proteins and endosomal uptake receptors more effectively than the hydroxyl/carbonyl-dominated surface of BB HT CQDs^38^. Second, the higher fluorescence quantum yield of BB MW CQDs amplifies the detectable intracellular signal at equivalent particle doses, contributing to the apparent superiority in fluorescence-based uptake quantification. Distinguishing between these two factors true variations in nanoparticle internalisation and differences in fluorescence efficiency per particle would require complementary analytical approaches, such as inductively coupled plasma mass spectrometry (ICP-MS) for quantitative intracellular carbon analysis, which remains an important direction for future investigations.

Taken together, the cellular uptake data reveal a passive, endocytic-activity-driven selectivity for cancer over non-cancerous cells that operates independently of any active targeting ligand^36^. Combined with the confirmed in vivo bioimaging performance in zebrafish larvae (Section 3.7) and the established cytocompatibility of both formulations (Section 3.5), these findings position BB MW CQDs as a compelling fluorescence imaging agent with inherent tumour-cell preferential accumulation.

## 4. Conclusions and Future Directions

Bearberry (Arctostaphylos uva-ursi) extract was used as a phytochemically rich, green precursor for the synthesis of fluorescent nitrogen- and oxygen-co-doped carbon quantum dots. Two routes were employed: hydrothermal (BB-HT CQDs) and microwave-assisted (BB-MW CQDs), both based on a common precursor system comprising citric acid, urea, and PEG 2000. A thorough analysis using UV-visible spectroscopy, photoluminescence, FTIR, XPS, and DLS shows that the two preparations’ physicochemical characteristics clearly differ from one another.. Hydrothermal treatment at 160 °C for 6 h yields CQDs with a more oxidised surface. These particles are enriched in hydroxyl, amine, and carbonyl functionalities. They also exhibit a higher fraction of graphitic carbon and a smaller hydrodynamic diameter (~7.13 nm). In contrast, microwave synthesis produces CQDs with higher surface carboxylate density and a less graphitised core. The hydrodynamic diameter of these particles is slightly greater (~9.65 nm). Zeta potential values in both systems are close to neutral (~0 mV).

Different functional outcomes are directly correlated with structural differences. The emission maxima of BB-HT CQDs is bathochromically shifted (478 nm vs. 467 nm).They also exhibit a more pronounced n-π* absorption feature. In parallel, they display a substantially lower DPPH inhibition. These trends are consistent with their higher surface density of hydroxyl and carbonyl groups. Such functionalities promote both hydrogen-atom transfer and electron-transfer pathways for radical scavenging. In contrast, BB-MW CQDs demonstrate superior optical efficiency. They exhibit a higher fluorescence quantum yield and enhanced cellular uptake. These properties are consistent with their increased surface carboxylate density and reduced non-radiative defect sites. BB-MW CQDs are hence more appropriate for imaging applications. According to in vivo findings, zebrafish larvae exhibit considerable absorption starting at 100 µg mL□^1^ (p < 0.05), with stronger accumulation at 250 and 500 µg mL□^1^ (p < 0.01).

Different functional results result directly from the structural disparities. The emission maxima of BB-HT CQDs is bathochromically displaced (478 nm vs 467 nm). Confocal fluorescence microscopy confirms intracellular uptake in both cell lines. BB-MW CQDs produce a statistically significant increase in fluorescence in RPE-1 cells at 200 µg mL□^1^ (p < 0.001). In HeLa cells, both formulations show consistent and substantial uptake at all tested concentrations. The higher accumulation in HeLa relative to RPE-1 cells suggests a degree of passive selectivity toward cancer cells. This behaviour may be advantageous for theranostic applications. Ongoing work will extend the physicochemical and biological characterisation. Future efforts will focus on targeted delivery strategies that exploit differences in surface chemistry between the two CQD types. Additional directions include combining phytochemical-derived antioxidant activity with photoluminescent imaging functionality, and validation in in vivo tumour models for theranostic applications.

ICP-MS quantification of intracellular carbon is the most pressing next step. Confocal fluorescence intensity conflates particle number with quantum yield BB MW CQDs emit more brightly per particle, so the uptake advantage over BB HT CQDs seen in Figure 15 may be partly optical rather than purely pharmacokinetic. Separating these contributions matters before any therapeutic dose rationale is built on these data. The endocytic pathway driving uptake is unresolved. Carboxylate-rich surfaces are typically internalised via clathrin-coated pits, but macropinocytosis cannot be excluded at the concentrations used here. Chlorpromazine and amiloride inhibition assays would settle this cleanly and determine whether the O–C=O surface enrichment of BB MW CQDs (19.06%) is mechanistically causal or merely correlative.

The passive HeLa/RPE-1 selectivity ratio 2.8-fold at 300 µg/mL for BB MW CQDs is large enough to build an active-targeting strategy on top of. EDC/NHS conjugation of folic acid to the carboxylate surface is chemically direct and would test whether receptor-mediated delivery amplifies this margin further. For BB HT CQDs, the graphitic core and surface hydroxyl groups are geometrically suited to doxorubicin loading via π-stacking; the particle’s own antioxidant activity could then offset drug-induced oxidative damage in bystander tissue a feature absent from conventional carbon nanocarriers.

BB MW CQDs at 467 nm emission and higher QY are well-positioned for two-photon excitation using 900–950 nm NIR laser sources. This would extend imaging depth from the ~200 µm accessible in the zebrafish larva to millimetre-scale penetration in mammalian tissue. Subcutaneous xenograft models in mice would be a logical next step after larval zebrafish, particularly given the passive tumour accumulation already demonstrated in HeLa cells.

Finally, the microwave route produces CQDs in minutes without pressurised equipment. Hydrothermal synthesis does not. For any scale-up programme, MW is the obvious starting point but conversion yield, batch-to-batch reproducibility, and lyophilisation energy have not been reported here. These are not trivial considerations for a green synthesis claim and should be quantified before pre-clinical scale-up is attempted.

## Declarations

None

## Acknowledgements

The authors express their gratitude to every member of the D.B. study group for their helpful criticism and thorough evaluation of the work. We sincerely thank the Indian Institute of Technology Gandhinagar for its financial and infrastructure support. The Central Instrumentation Facility (CIF) at IIT Gandhinagar is acknowledged by the authors for granting access to the characterisation tools utilised in this investigation, including AFM, FTIR, XRD. A.S. thanks financial support from the Prime Minister’s Research Fellowship and the Government of India’s Ministry of Education. The Indian National Young Academy of Sciences acknowledges D.B. as a member. J.P and S.B. thanks IIT Gandhinagar’s Department of Biological Sciences and Engineering for providing the resources and research environment needed to complete this work. The work in the host labs is supported by funding from ANRF-CRG, GSBTM, MoES-STARS, CCRH, IITGN.

## Author Contributions

D.B.: Funding acquisition, writing, review and editing, supervision, resources, and conceptualisation. S.B, J.P., B.P., A.S: Conceptualisation, research, formal analysis, methodology, first draft writing, and visualisation. S.B.: Research, Approach. AKM and A.S.: Data curation, methodology, and investigation.

## References

1. Daby, T. P. M., Modi, U., Yadav, A. K., Bhatia, D. & Solanki, R. Bioimaging and therapeutic applications of multifunctional carbon quantum dots: Recent progress and challenges. Nanotechnol. 8, 100158 (2025).

2. Kumar, A. et al. Clinical Applications of Targeted Nanomaterials. Pharmaceutics 17, 379 (2025).

3. Functionalized Nanomaterials for Detecting Environmental Pollutants - Dave - 2024 - ChemistrySelect - Wiley Online Library. https://chemistry-europe.onlinelibrary.wiley.com/doi/full/10.1002/slct.202402477.

4. Barve, K., Singh, U., Yadav, P. & Bhatia, D. Carbon-based designer and programmable fluorescent quantum dots for targeted biological and biomedical applications. Mater. Chem. Front. 7, 1781–1802 (2023).

5. Yadav, P. et al. Green emitting carbon quantum dots (GCQDs) to probe endocytic pathways in cells; for tissue and in vivo bioimaging. 2022.04.23.489248 Preprint at 10.1101/2022.04.23.489248 (2022).

6. Benner, D., Yadav, P. & Bhatia, D. Red emitting carbon dots: surface modifications and bioapplications. Nanoscale Adv. 5, 4337–4353 (2023).

7. Kumar, A. et al. Selective cellular uptake and cytotoxicity effects of fluorescent carbon dots: a comparative study in cancer and normal cells. Mater. Adv. 7, 495–508 (2026).

8. Singh, U. et al. Water stable, red emitting, carbon nanoparticles stimulate 3D cell invasion via clathrin-mediated endocytic uptake. Nanoscale Adv. 4, 1375–1386 (2022).

9. Next□Generation Cancer Theragnostic: Applications of Carbon Quantum Dots - Kumar - 2025 - ChemNanoMat - Wiley Online Library. https://aces.onlinelibrary.wiley.com/doi/10.1002/cnma.202500061.

10. Barot, R. B. et al. Novel carbon nanoparticles derived from Bougainvillea modulate vegetative growth via auxin–cytokinin signaling in Arabidopsis. Chem. Pap. 78, 4733–4750 (2024).

11. Bhatia, D. et al. Quantum dot-loaded monofunctionalized DNA icosahedra for single-particle tracking of endocytic pathways. Nat. Nanotechnol. 11, 1112–1119 (2016).

12. Sugier, P. et al. Chemical Characteristics and Antioxidant Activity of Arctostaphylos uva-ursi L. Spreng. at the Southern Border of the Geographical Range of the Species in Europe. Molecules 26, 7692 (2021).

13. Barve, K. et al. Red fluorescent carbon nanoparticles derived from Spinacia oleracea L.: a versatile tool for bioimaging and biomedical applications. Mater. Adv. 4, 6277–6285 (2023).

14. Singh, R. K., Kumar, R., Singh, D. P., Savu, R. & Moshkalev, S. A. Progress in microwave-assisted synthesis of quantum dots (graphene/carbon/semiconducting) for bioapplications: a review. Mater. Today Chem. 12, 282–314 (2019).

15. Libra, J. A. et al. Hydrothermal carbonization of biomass residuals: a comparative review of the chemistry, processes and applications of wet and dry pyrolysis. Biofuels 2, 71–106 (2011).

16. Qi, H. et al. Novel N-doped carbon dots derived from citric acid and urea: fluorescent sensing for determination of metronidazole and cytotoxicity studies. RSC Adv. 13, 2663–2671.

17. Sun, Y.-P. et al. Quantum-Sized Carbon Dots for Bright and Colorful Photoluminescence. J. Am. Chem. Soc. 128, 7756–7757 (2006).

18. Hola, K. et al. Carbon dots—Emerging light emitters for bioimaging, cancer therapy and optoelectronics. Nano Today 9, 590–603 (2014).

19. Wang, D. et al. Carbon dots as new antioxidants: Synthesis, activity, mechanism and application in the food industry. Food Chem. 475, 143377 (2025).

20. Ding, H., Yu, S.-B.Wei, J.-S. & Xiong, H.-M. Full-Color Light-Emitting Carbon Dots with a Surface-State-Controlled Luminescence Mechanism. ACS Nano 10, 484–491 (2016).

21. Lim, S. Y., Shen, W. & Gao, Z. Carbon quantum dots and their applications. Chem. Soc. Rev. 44, 362– 381 (2014).

22. Liu, M. L., Chen, B. B., Li, C. M. & Huang, C. Z. Carbon dots: synthesis, formation mechanism, fluorescence origin and sensing applications. Green Chem. 21, 449–471 (2019).

23. Jain, S., Sahu, N., Bhatia, D. & Yadav, P. Cellular uptake and viability switching in the properties of lipid coated carbon quantum dots for potential bioimaging and therapeutics. Nanoscale Adv. 6, 5069–5079 (2024).

24. Harris, I. S. et al. Glutathione and thioredoxin antioxidant pathways synergize to drive cancer initiation and progression. Cancer Cell 27, 211–222 (2015).

25. Rodríguez-Varillas, S., Espina-Casado, J., Badía Laíño, R., Fernández-González, A. & Fontanil López, T. Carbon dots in oncology: multifunctional nanoplatforms for diagnosis, targeted therapy, and drug discovery. Drug Discov. Today 30, 104470 (2025).

26. Electrochemical Study of DPPH Radical Scavenging for Evaluating the Antioxidant Capacity of Carbon Nanodots | The Journal of Physical Chemistry C. https://pubs.acs.org/doi/10.1021/acs.jpcc.7b05353.

27. Salvia Miltiorrhiza-Derived Carbon Dots as Scavengers of Reactive Oxygen Species for Reducing Oxidative Damage of Plants | ACS Applied Nano Materials. https://pubs.acs.org/doi/10.1021/acsanm.0c02419.

28. Yadav, P. et al. Dopamine-Functionalized, Red Carbon Quantum Dots for In Vivo Bioimaging, Cancer Therapeutics, and Neuronal Differentiation. ACS Appl. Bio Mater. 7, 3915–3931 (2024).

29. In vivo characterization of hair and skin derived carbon quantum dots with high quantum yield as long-term bioprobes in zebrafish | Scientific Reports. https://www.nature.com/articles/srep37860.

30. Carbon Quantum Dots for Zebrafish Fluorescence Imaging | Scientific Reports. https://www.nature.com/articles/srep11835.

31. Sun, Y.-P. et al. Quantum-sized carbon dots for bright and colorful photoluminescence. J. Am. Chem. Soc. 128, 7756–7757 (2006).

32. Wang, Y. et al. Enhanced dispersion stability of gold nanoparticles by the physisorption of cyclic poly(ethylene glycol). Nat. Commun. 11, 6089 (2020).

33. Bian, Z. et al. Bottom-up carbon dots: purification, single-particle dynamics, and electronic structure. Chem. Sci. 16, 4195–4212 (2025).

34. Qu, D. & Sun, Z. The formation mechanism and fluorophores of carbon dots synthesized via a bottom-up route. Mater. Chem. Front. 4, 400–420 (2020).

35. Recouvreux, M. V. & Commisso, C. Macropinocytosis: A Metabolic Adaptation to Nutrient Stress in Cancer. Front. Endocrinol. 8, 261 (2017).

36. Yadav, P. et al. Tissue-Derived Primary Cell Type Dictates the Endocytic Uptake Route of Carbon Quantum Dots and In Vivo Uptake. ACS Appl. Bio Mater. 6, 1629–1638 (2023).

37. Selective cellular uptake and cytotoxicity effects of fluorescent carbon dots: a comparative study in cancer and normal cells - Materials Advances (RSC Publishing) DOI:10.1039/D5MA00781J. https://pubs.rsc.org/en/content/articlehtml/2026/ma/d5ma00781j.

38. Nanoparticle Surface Functionality Dictates Cellular and Systemic Toxicity | Chemistry of Materials. https://pubs.acs.org/doi/10.1021/acs.chemmater.7b01979.

